# Small-molecule binding and sensing with a designed protein family

**DOI:** 10.1101/2023.11.01.565201

**Authors:** Gyu Rie Lee, Samuel J. Pellock, Christoffer Norn, Doug Tischer, Justas Dauparas, Ivan Anischenko, Jaron A. M. Mercer, Alex Kang, Asim Bera, Hannah Nguyen, Inna Goreshnik, Dionne Vafeados, Nicole Roullier, Hannah L. Han, Brian Coventry, Hugh K. Haddox, David R. Liu, Andy Hsien-Wei Yeh, David Baker

**Author notes:** Equal contribution (Gyu Rie Lee, Samuel J. Pellock, Christoffer Norn). Co-corresponding authors. (D.B.); (A.H.W.Y.).

## Abstract

Despite transformative advances in protein design with deep learning, the design of small-molecule–binding proteins and sensors for arbitrary ligands remains a grand challenge. Here we combine deep learning and physics-based methods to generate a family of proteins with diverse and designable pocket geometries, which we employ to computationally design binders for six chemically and structurally distinct small-molecule targets. Biophysical characterization of the designed binders revealed nanomolar to low micromolar binding affinities and atomic-level design accuracy. The bound ligands are exposed at one edge of the binding pocket, enabling the *de novo* design of chemically induced dimerization (CID) systems; we take advantage of this to create a biosensor with nanomolar sensitivity for cortisol. Our approach provides a general method to design proteins that bind and sense small molecules for a wide range of analytical, environmental, and biomedical applications.

## Main

The design of small-molecule–binding proteins with high affinity and specificity is of considerable current interest. For example, biosensors and switches that undergo dimerization upon ligand binding (chemically-induced dimerization (CID)) are broadly useful, but most approaches to develop new inducible systems focus on the discovery and engineering of natural proteins^1,2,3^, as general methods are not currently available for designing protein-small molecule interactions that drive protein association. Previous efforts to design small-molecule–binding proteins have employed physics-based methods to engineer natural proteins^2–6^ or design small-molecule binders *de novo^7–9^*, but these approaches have yielded binders only for a single small-molecule target, perhaps because low scaffold diversity has limited the molecular space that can be recognized. Large libraries of de novo protein scaffolds have been generated to address this^10,11^, but sequence design methods that can satisfy the structural diversity of a large scaffold set while also designing precise small-molecule interactions are lacking. Thus, a more general method is needed that samples backbone space broadly and encodes it accurately to generate a diverse set of small-molecule binding proteins.

We hypothesized that a more general solution to the problem of small-molecule binder design could be attained by combining advances in deep learning-based protein fold generation and sequence design. For the former, we reasoned that a large set of scaffolds housing stable pockets could enable design of binding sites for a wide variety of small molecules, and that the most suitable folds would be both compact (to keep the designs small and modular) and diversifiable (to enable generation of a wide variety of binding sites). For downstream CID applications, we sought a structural solution with the bound ligand sufficiently exposed to enable modulation of a designed protein interaction by ligand binding. Based on these criteria, we chose the compact but readily diversifiable NTF2 fold: despite their relatively small size (∼120 residues), natural family members can bind small molecules, the fold has already been shown to be compatible with small molecule binder design^4,12^, and the bound ligands are exposed at one end of the binding pocket. For sequence design, we reasoned that the recently developed LigandMPNN, a deep learning model for protein sequence design trained on protein-small molecule complexes, could generate protein-ligand interactions more effectively than previous approaches that struggle to balance protein-ligand and intramolecular protein-protein interactions simultaneously^12^.

### Computational design of small-molecule–binding proteins

The NTF2 fold is composed of 3 helices and a curved, 6-stranded β-sheet, which together form the large internal pocket characteristic of this fold family (**Fig. 1A**). The natural diversity of this fold is mediated by long irregular loops and unique quaternary structures, both of which affect pocket geometry and function^13,14^. We set out to design a family of NTF2s with diverse pocket geometries to accommodate a wide range of small molecules, but that are strictly monomers with only minimal loops to maintain their modularity and designability. To achieve this, we generated NTF2s with trRosetta-based family-wide hallucination^15^ (Set 1: 1,615 scaffolds), redesigned these backbones with ProteinMPNN^16^ and selected those that fold to the designed structure with AlphaFold^17^ (Set 2: 3,230 backbones), and in addition parametrically generated backbones with Rosetta^10^, redesigned these with ProteinMPNN, and validated their structure with AlphaFold (Set 3: 6,838 backbones) (**Fig. S1**).

**Figure 1.**
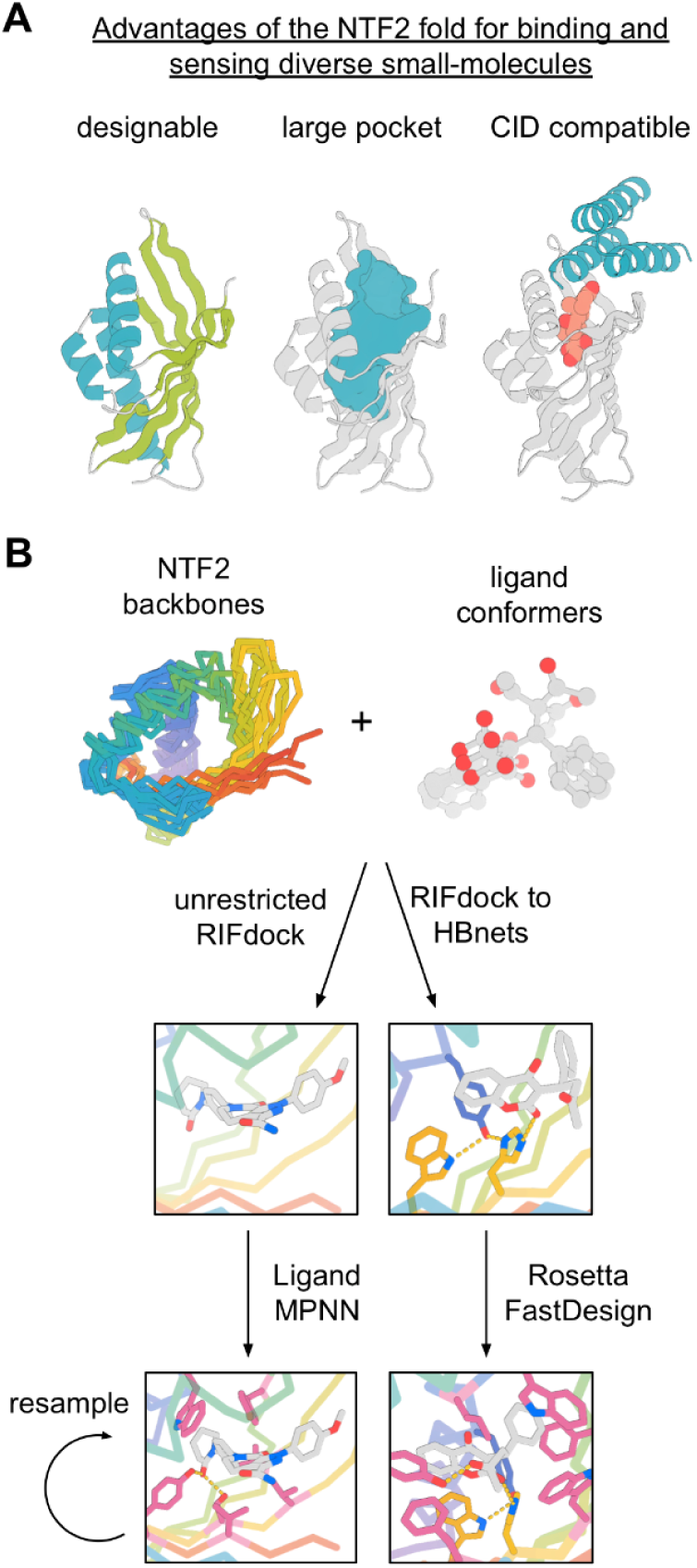
Rationale and design pipeline for binding and sensing small-molecules in the NTF2 fold. (**A**) The NTF2 fold has a designable fold made up of ideal secondary structure elements (left panel, helices in teal and strands in green), the large pocket (teal, middle panel) of NTF2s can bind various small-molecules, and the binding mode of small-molecules in NTF2s enables the generation of CIDs (right panel, NTF2 in gray, CID agent in teal, and small-molecule in red). (**B**) Design pipeline for small-molecule–binding in the NTF2 fold.

With a designed family of over 10,000 NTF2s with diverse pocket geometries in hand, we used RIFdock^7^ to place six chemically and structurally distinct small-molecules in the central pocket of these backbones (**Fig. 1B**), including the stress hormone cortisol (HCY)^18^, the anticoagulant warfarin (WRF)^19^, the muscle relaxant rocuronium (ROC)^20^, the anticoagulant apixaban (APX)^21^, the active metabolite of the anticancer drug irinotecan (SN-38)^22^, and the hormone 17-α-hydroxyprogesterone (OHP)^23^ (**Fig. 2A**). Designing polar interfaces remains an outstanding challenge in protein design^24,25^, especially for small-molecule binding proteins, where an internal pocket with polar residues is needed to interact with polar functional groups of a ligand without destabilizing the protein fold. By explicitly docking polar functional groups of small molecules to pre-installed hydrogen bond networks (HBNets) present in Set 1 scaffolds (approach 1) or leveraging deep learning-based design methods trained on protein-small molecule interactions (approach 2), we reasoned that we could generate highly preorganized polar contacts while maintaining the integrity of the protein fold. For approach 1, we docked HCY, WRF, ROC, APX, and SN-38 into Set 1 backbones and required that at least one protein-small-molecule interaction was mediated by an HBNet residue, after which we used native-sequence guided Rosetta design as previously described^15^. For approach 2, we used RIFdock without constraints to place OHP, APX, and SN-38 into scaffold Sets 2 and 3 and performed sequence design with LigandMPNN, a version of ProteinMPNN trained on small molecule-protein complexes that explicitly considers the ligand during sequence design. To select amongst the resulting designs from both approaches, we used Rosetta to calculate the number of hydrogen bonds between the protein and ligand, the binding energy (ddG), and contact molecular surface (CMS)^26^ (**Fig. S2A**). For design approach 2, we also selected designs based on recapitulation of the fold and binding site by single-sequence AlphaFold predictions (**Fig. S2B**). After filtering, we obtained oligonucleotides encoding the binders for experimental characterization, including 630 for HCY, 1,661 for ROC, 16,276 for WRF, 9,024 for APX, 19,390 for SN-38, and 7,573 for OHP.

**Figure 2.**
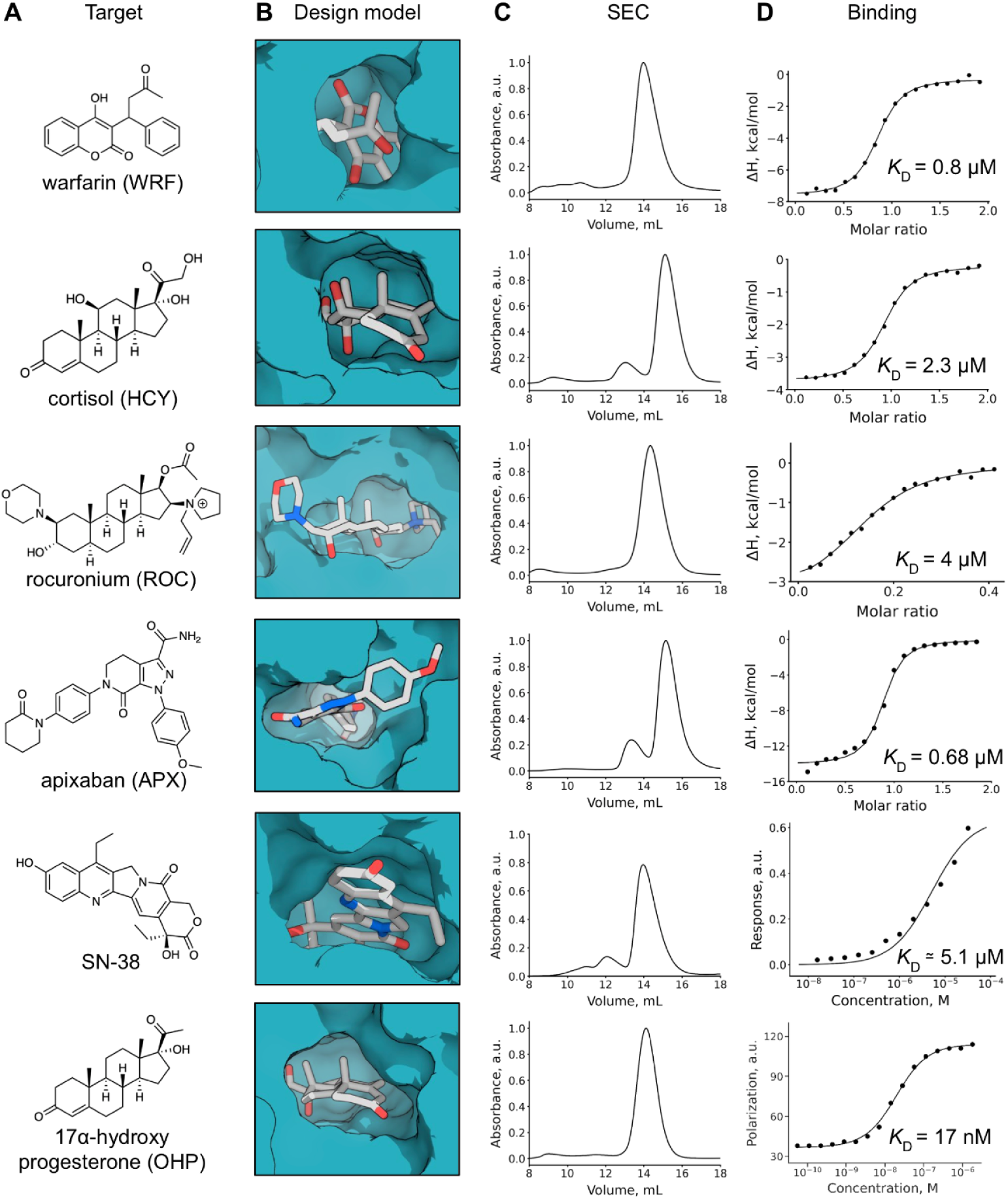
Characterization of small-molecule–binding proteins. (**A**) Chemical structures of small-molecule targets. (**B**) Surface representation (teal) of designed binders in complex with small-molecule targets (gray) (**C**) Size-exclusion chromatography traces of designed binders. (**D**) ITC binding isotherms of designed proteins titrated with cognate small molecule targets.

### Characterization of small-molecule binding proteins

Designs were ordered as synthetic oligonucleotides and transformed into yeast. Binding screens were performed with yeast surface display coupled to fluorescence-activated cell sorting (FACS), where interaction with a biotinylated small-molecule target enables labeling with streptavidin-phycoerythrin (SAPE). Binding signal was detected for all six targets and significant enrichment was observed after multiple rounds of sorting (**Fig. S3**). Deep sequencing of the final sorted populations revealed 1, 46, 19, 8, 8, and 117 unique hits for HCY, WRF, ROC, APX, SN-38, and OHP, respectively. AF2 and Rosetta metrics for the identified hits revealed high confidence (pLDDTs ranging from 86.0 to 95.0), accuracy (Cα RMSDs less than 2.0 Å), and extensive molecular interactions (target median ddG < −30) and high shape complementarity (target median CMS > 240) to the ligand . and close agreement with AlphaFold models, suggesting high accuracy of the design method (**S4A,B**).

We performed site saturation mutagenesis (SSM) experiments on a selected set of binders identified from yeast display screening to assess the sequence-function relationship and confirm the designed binding mode. Overall, analysis of the SSMs revealed high conservation of the designed protein-ligand interactions, including key hydrogen bond and hydrophobic interactions (**Fig. S6**). Notably, contacts from pre-installed HBNets were highly conserved in the SSM. Furthermore, key residues that preorganize these residues were also conserved. Design approach 2 based on LigandMPNN was also able to design pi-pi or CH-pi interactions, which are more complex for physically based force fields to accurately model^27^.

Select designs from this set of putative binders were expressed in *E. coli* and purified to assess their solubility, oligomeric state, and binding affinity to their cognate targets. All selected designs displayed some level of expression and 23 of 33 were distinct monomers or dimers by size-exclusion chromatography (SEC) (**Fig. 2C, Fig. S5**). The binding affinities of the purified proteins for their target small-molecules were determined by isothermal titration calorimetry (ITC), fluorescence polarization, and bio-layer interferometry, and the *K*_D_ of the highest affinity binders across all targets ranged from micromolar to low nanomolar (**Fig. 2D**). For rocuronium, the most enriched hit identified by yeast display showed poor solution properties by SEC (**Fig. S5**), so we optimized this sequence by combining favorable mutants from the SSM and a mutation based on an AlphaFold multimer prediction that showed the formation of a dimer that likely precludes binding to rocuronium (**Fig. S6 and S7**). Combining these mutations improved solubility and enabled characterization of binding (**Fig. 2**). We also optimized the most highly enriched hit for SN-38 by incorporating three substitutions identified in an SSM experiment which enabled measurement of binding.

### Structural characterization of designed cortisol-binding protein

To further assess the accuracy of designed small-molecule binding proteins, experimental structures of protein-ligand complexes were determined. To aid in crystallization, ProteinMPNN was used to redesign the surface of select designs given its previous success in yielding crystallizable sequences^28^. We crystallized a proteinMPNN-redesign of the cortisol-binder hcy129, and obtained the 1.5 Å crystal structure of this design in complex with cortisol (**Fig. 3A and S8**). Structural alignment of the crystal structure to the design model revealed a Cα RMSD of 1.1 Å over 129 residues (**Fig. 3A**). In addition to the overall accuracy of the fold, key hydrogen bonding residues, such as W17, Y18, and Y108, and the binding orientation of cortisol closely match the design model (**Fig. 3C,D**). Notably, multiple polar interactions are mediated by a pre-installed HBNet unique to scaffold Set 1, demonstrating that this approach worked as intended to generate preorganized polar contacts.

**Figure 3.**
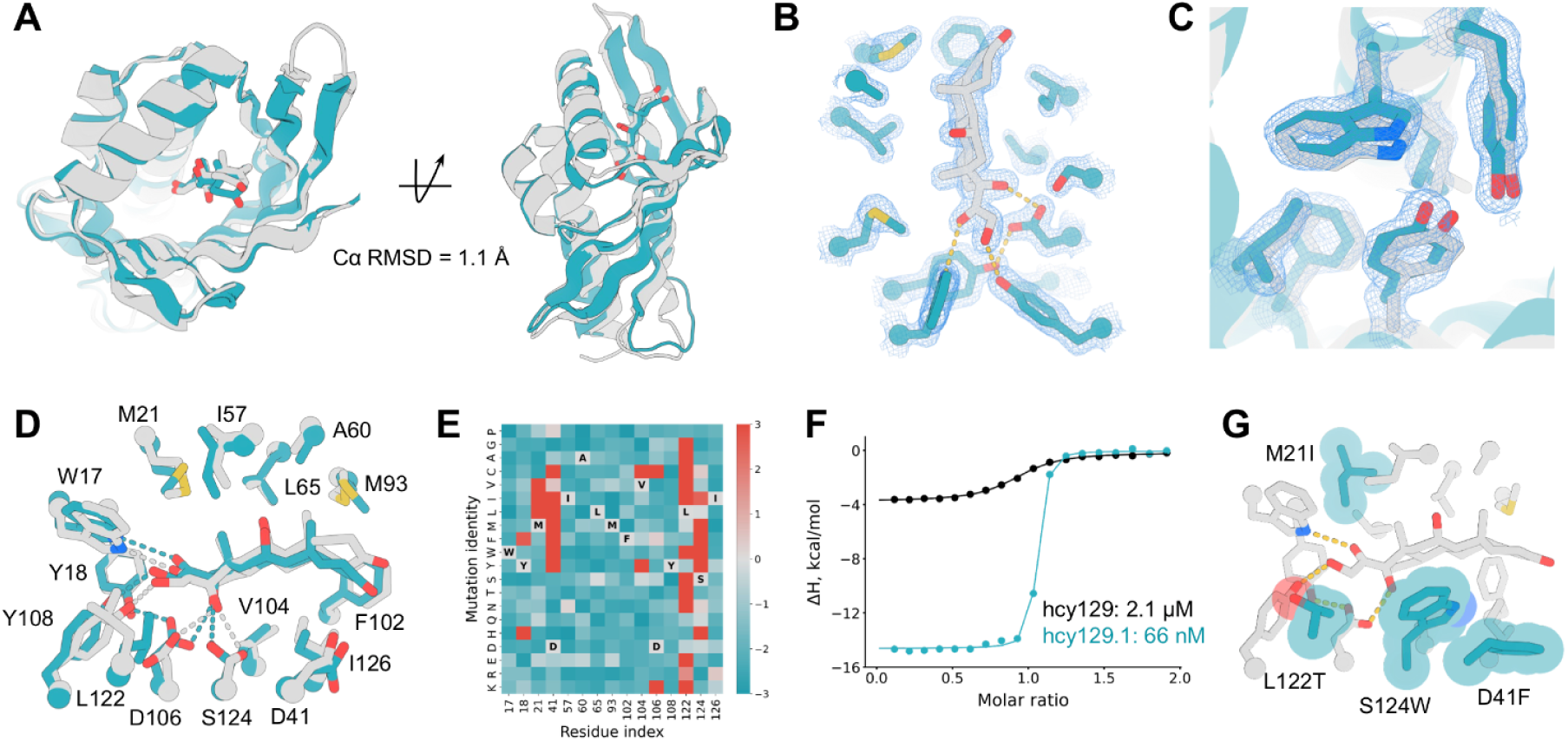
Structural characterization and affinity maturation of a designed cortisol-binding protein. (**A**) Structural superposition of design model (gray) and crystal structure (teal) of hcy129_mpnn5. (**B**) Zoom-in of binding site of hcy129_mpnn5 with 2Fo-Fc density contoured at 1.5 σ. (**C**) Overlay of design model (gray) and crystal structure (teal) zoomed into the folded core of hcy129_mpnn5. (**D**) Zoom-in overlay of design model (gray) and crystal structure (teal) binding sites. (**E**) SSM heatmap of binding residues of hcy129 (teal: weaker binding, gray: no change, red: tighter binding). (**F**) ITC binding isotherms of hcy129 and hcy129.1. (**G**) Zoom-in of the binding site of an AlphaFold-predicted structure of hcy129.1 with cortisol docked using GALigandDock and mutated residues show in teal.

### Design and characterization of a cortisol-induced heterodimer

To improve the binding affinity of hcy129 for use as a cortisol biosensor, which is typically at low nanomolar concentrations in human saliva and blood^29^, we screened a library of combinatorial mutants based on favorable mutations identified from the SSM of hcy129 (**Fig. 3E**) by yeast display, and observed significant improvements in binding affinity (**Fig. S9**). We selected the best variant from this library for production in *E. coli* and characterization by ITC. This variant, hcy129.1, displayed a K_D_ of 66 nM, a 35-fold improvement in affinity compared to the original design (**Fig. 3F**). Docking cortisol into the AlphaFold model of hcy129.1 with GALigandDock^30^ revealed that improvements in affinity are likely a result of improved hydrophobic interactions with cortisol (**Fig. 3G**). With a binder that recognizes cortisol at physiologically relevant concentrations in hand, we set out to design a cortisol-dependent dimerization system. The binding mode of cortisol in the NTF2 fold leaves part of its structure exposed, enabling the design of proteins that form a ternary complex that interfaces with the ligand-bound state of the NTF2 (**Fig. 4A**). First, the surface at the opening of the pocket of hcy129.1 was redesigned to facilitate docking and design of protein binders to the hcy129.1-cortisol complex. After generating this new variant, hcy129.1_CID, we performed RIFdock against the hcy129.1_CID-cortisol complex with a previously described miniprotein scaffold library^26^ (**Fig. 4A**). This approach generated numerous ternary complexes where both the miniprotein and hcy129.1 interact with each other and cortisol (**Fig. 4A**). Sequence design of the resulting complexes were carried out with both Rosetta FastDesign and ProteinMPNN, and the heterodimeric complexes were validated with AlphaFold2 and filtered for pAE < 10 and pLDDT > 85.

**Figure 4.**
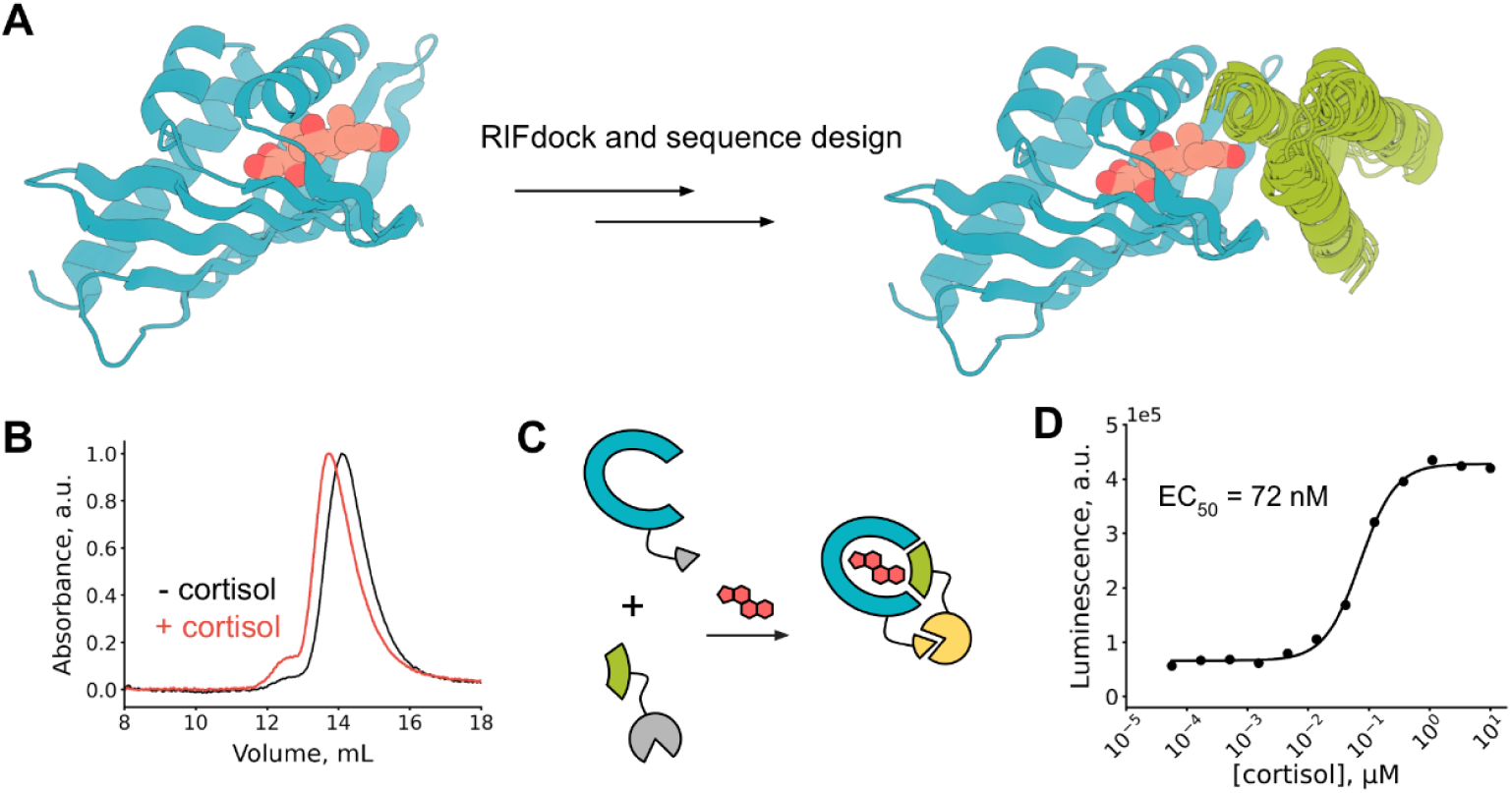
Design and characterization of a cortisol-dependent heterodimer. (**A**) Design pipeline for cortisol-induced heterodimerization. Starting from a model of hcy129.1_CID (teal) in complex with cortisol (peach) (left), we used RIFdock and a library of designed 3-helix bundles (green) to generate thousands of putative ternary complexes and the amino acid sequences encoding these complexes were generated with FastDesign and ProteinMPNN (right).(**B**) Size-exclusion chromatography traces of an equimolar solution of hcy129.1 and miniH11 (1 μM) in the presence (red) or absence (black) of cortisol (10 μM). (**C**) Schematic diagram of the designed cortisol-induced heterodimerization coupled with a binary split luciferase to create a biosensor. (**D**) Cortisol dependent luminescent responses in an equimolar solution of hcy129.1_CID-SmBiT and mini11_LgBiT (200 nM), showing sensitivity at low nM range.

The designed minibinders were displayed on yeast and binding to biotinylated hcy129.1_CID was assessed in the presence or absence of cortisol by FACS (**Fig. S10**). Populations enriched for binding to hcy129.1_CID in the presence of cortisol were collected and colonies were picked to identify putative CIDs. We expressed and purified a select subset of these and characterized them *in vitro* to confirm cortisol-induced dimerization. As an initial test, we purified and combined both hcy129.1_CID and the minibinders in the presence or absence of cortisol and analyzed them by SEC to detect a change in elution volume. For 9 out of 12 designs tested, we observed a clear shift by SEC towards a higher molecular weight species in the presence of cortisol (**Fig. 4B and Fig. S11**). Of these, we analyzed one, miniH11, by native mass spectrometry, which revealed a molecular weight for the hcy129.1_CID-cortisol-miniH11 ternary complex (**Fig. S12A**). Together, these data suggest that the designed proteins form a cortisol-dependent CID.

As a proof-of-concept for cortisol sensing based on the designed CID, we genetically fused hcy129.1_CID and miniH11 to the SmBiT and LgBiT components of the NanoBiT system, respectively, which reconstitutes the NanoBiT luciferase and generates luminescence when brought in close proximity by a molecular interaction^31^ (**Fig. 4C**). We expressed and purified the CID-NanoBiT fusion constructs and titrated them with cortisol, which generated a luminescent signal with an estimated EC_50_ of 72 nM (**Fig. 4D**). This closely matches the *K*_D_ of hcy129.1 for cortisol, which suggests that a specific interaction between hcy129.1_CID and cortisol promotes association of miniH11. To assess the binding affinity of the CID components in the absence of cortisol, we titrated miniH11-LgBiT with increasing concentrations of hcy129.1_CID-SmBiT, which revealed an estimated K_D_ of ∼5 μM (**Fig. S12B**), approximately 2 orders of magnitude greater than the EC_50_ identified for cortisol-induced dimerization, indicating that the dimerization observed at low concentrations of the CID pair is dependent on the presence of cortisol. Taken together, these data demonstrate that the NTF2-based small-molecule binders designed in this study can be readily engineered to serve as sensors for small-molecules of interest.

## Conclusion

By integrating the deep learning methods trRosetta hallucination for backbone generation, ProteinMPNN and LigandMPNN for sequence design, and AlphaFold for filtering, we show that scaffold sets around a fold of interest that sample novel sequence and structure space can be readily generated with considerable utility for the design of functional proteins, as demonstrated by the design of binders to six structurally distinct targets. We show that the NTF2 fold not only can harbor a wide range of binding pockets, but also enables the modular construction of CID systems, as illustrated by the cortisol biosensor. The computational approach developed here provides a general approach not only for the design of small-molecule–binding proteins but also for the creation of protein heterodimers that respond to a selected small-molecule, broadening the toolbox for programmable synthetic biology and a diverse array of biomedical applications.

## Methods

### Generation of NTF2 scaffolds

Set 1 backbones were generated as previously described^15^. For set 2 and set 3 backbones, hallucinated^15^ and algorithmically generated^10^ backbones were used as input for ProteinMPNN. For hallucinated backbones, two separate ProteinMPNN design approaches were taken, one in which native HBNet residues were kept fixed and one in which the entire protein was allowed to be redesigned, and both design runs were performed with and without polar residue-biased weights on amino acid (aa) compositions. The algorithmically generated backbones were also designed with and without a polar-aa bias term applied. Structures of resulting sequences were predicted by AlphaFold2 in single-sequence mode, using model 4 with 10 recycles, and structures with Cα RMSDs less than 1 Å compared to the original scaffold and an average plDDT greater than 92 were finally chosen.

### Generation of ligand conformers

For ROC, HCY, WRF, APX, and SN-38, 2D chemical structures were made in ChemDraw and exported as SDfiles or SMILES strings, which served as input for RDKit to generate initial 3D models of the target structures with the ETKDGv2 method^32^. For the first set of designs based on set1 backbones, partial charges were assigned using antechamber^33^ and geometry optimization was performed using xtb^34^. Additional sampling of ligand conformers was performed with the CSD conformer generator^35^ and final sets of 15 conformers of fewer were selected manually as input for RIFgen. 200 conformers of OHP, APX, SN-38 were generated for each ligand in design approach 2 and the structures were optimized using the MMFF force field^36^. The conformers were clustered based on pairwise RMSD. 1, 26, and 4 cluster representatives with lowest energy were selected respectively for OHP, APX, SN-38, and final minimization of the ligand conformations was done by using the AIMnet potential^37^.

### Rosetta ligand parameter generation

Conformers generated for each target small-molecule were converted into MOL2 format using openbabel and used as inputs to generate Rosetta parameter files. These parameters were provided when using the generic potential to perform Rosetta calculations^30^.

### Computational design of small-molecule-binding proteins

Backbone generation, ligand docking, and sequence design for set 1 backbones was performed as described previously^15^. Backbones were generated with the TrRosetta-based hallucination method using constraints derived from NTF2-like structures and homology models including sequence and geometric constraints for recapitulation of native hydrogen-bonding networks^15^. These hydrogen-bonding networks (HbNet) were utilized for docking the polar functional groups of the small-molecules. For each target small-molecule, functional groups that can serve as hydrogen bond donors or acceptors were identified, and biased RIFdock was performed for all possible combinations of ligand atom and HbNet residue pairs. Sequences of the RIFdock solutions of protein and ligand complexes were redesigned using Rosetta FastDesign^38^ with position-specific scoring matrix (PSSM) sequence constraints, while the amino acid identities of the residues forming hydrogen-bonds to the target ligand were kept fixed. The PSSM was constructed using the collected NTF2-like sequences. Rosetta metrics such as ddG, contact molecular surface (CMS), solvent accessible surface area between the ligand and the protein (SASA), and the number of hydrogen bonds (nHb) were used to select the designs to test (**Table S1**), and the sequences were finally clustered based on sequence identity.

For design approach 2, all conformers of OHP, APX, and SN-38 were docked to scaffolds Set 2 and 3 using unconstrained RIFdock. The docks were initially filtered based on their shape complementarity to the scaffold, estimated by the protein-ligand contact molecular surface area calculated using Rosetta^26^. The filtered docks underwent an initial round of sequence design using LigandMPNN followed by Rosetta FastRelax^39,40^. The backbone and the coordinates of the ligand atoms were used as input for LigandMPNN and no sequence constraints were applied at this stage. Eight sequences were generated with temperature T=0.2 per input, and 3 iterations of LigandMPNN and RosettaPackMin (before the last step) or RosettaFastRelax (last step) were performed. Distance restraints were applied for selected sets of ligand atoms and protein Cα atoms in the binding pocket to keep the ligand in place after sequence change. The designs were filtered using Rosetta metrics, including ddG, contact molecular surface (CMS), and the number of hydrogen bonds (nHb) formed between the protein and ligand. We used AlphaFold2 to validate the designed sequences, and filtered based on plDDT, Cα RMSD, and binding site residue side chain RMSD (**Table S1**). To increase the sequence diversity while maintaining structure-prediction quality of the designed sequences, GALigandDock was performed on the AF2 models of the selected designs, and the docking solutions with ligand all-atom RMSD lower than 2 Å were used as inputs for the next iterative round of sequence design. This process allowed sampling additional degrees of freedom of the ligand as ligand docking also optimized the internal torsion angles of the ligands. Second round of sequence design based on the same design scheme consisting of LigandMPNN and RosettaRelax was applied to the updated inputs, and the final set of designs to be ordered were selected using Rosetta and AF2 metrics (**Table S1**).

### Library assembly and yeast display screening

DNA encoding designed small-molecule–binding proteins were ordered as single-stranded synthetic oligos from Twist Bioscience and assembled as described previously^41^. The assembled library DNA was mixed with linearized pETCON3 plasmid and the resulting solution was transformed into competent EBY100 yeast by electroporation as previously described^42^. The yeast library was first sorted for expression, and any cells labeled by anti-c-Myc-fluorescein isothiocyanate (anti-c-Myc-FITC, Immunology Consultants Laboratory) were collected and then grown for 2-3 days in c -Trp -Ura selection media. To assess binding, the cells recovered from the expression sort were incubated with biotinylated ligand, anti-c-Myc-FITC, and streptavidin conjugated to phycoerythrin (SAPE, ThermoFisher) in PBSF (phosphate buffered saline, Fisher Scientific with 0.1% bovine serum albumin). Yeast cells labeled by both anti-c-Myc-FITC and SAPE were collected and this same process was repeated for at least two more rounds of sorting. The concentrations of the biotinylated ligand and SAPE were chosen differently for each sort (**Fig. S3**). Final cell populations were prepared for deep sequencing or streaked in c -Trp -Ura plates and individual colonies were sequenced by colony PCR and Sanger sequencing to identify functional sequences.

### Site saturation mutagenesis library preparation and sorting

For each putative binder amino acid sequence, single site positions were mutated to all possible amino acids and were ordered as synthetic oligos from Twist or Agilent. Oligos with mutations on the N terminal half side were assembled with a wild type sequence gblock ordered from Integrated DNA Technologies (IDT). The oligos and the gblock were constructed to have constant overlap for assembly. The same was applied to the oligos with mutations on the C terminal half, and after combining the two DNA pools, yeast cell transformation by electroporation and FACS was performed in the same way as described previously.

### Combinatorial mutagenesis library preparation and sorting

#### Cortisol binder hcy129

DNA fragments with combinatorial mutations were ordered as eblocks from IDT with BsaI restriction sites and pETCON3 vector overlapping sequences. Eblocks corresponding to N or C terminal ends of the sequence were pooled and were stitched together using Golden Gate assembly. The assembled library was chemically transformed^43^ to competent EBY100 yeast cells with the linearized pETCON3 vector.

#### Rocuronium binder roc22182

We ordered synthetic oligonucleotides including degenerate codons (opools, IDT) optimized with SwiftLib^44^ to construct a combinatorial mutagenesis library. Oligonucleotides were cloned into a pETCON3 vector and transformed into yeast cells using electroporation.

#### SN-38 binder iri807

The structures of all possible combinatorial mutant sequences were predicted using AlphaFold2, and 153 sequences that yielded predictions with plDDT > 90.0 and Cα RMSD < 1.5 Å were chosen. The DNA fragments of the selected sequences were ordered as eblocks, chemically transformed to yeast cells, and the binding of each clone to SN-38 were analyzed using the Attune flow cytometer (Thermo Fisher). The sequences with the best binding signals were tested for binding using BLI.

### Deep sequencing and analysis

The collected yeast cells after FACS were grown in c -Trp -Ura media with 2% glucose for 2 to 3 days, and 3e7 to 5e7 cells were used to extract the plasmids (Zymoprep, Zymo Research). Two rounds of qPCR amplifications were performed using the same protocol as the design library assembly for DNA amplification and attachment of Illumina and pool specific barcodes. The purified DNA samples were sequenced using Illumina MiSeq sequencing.

The sequencing outputs were downloaded in FASTQ format, and we used the program PEAR^45^ to merge the paired end reads. Reads matching the ordered design amino acid sequences were counted and used for analyzing the binding enrichments. For each sequence the frequencies were calculated for all sorts, and the log ratio of the binding frequency to the expression frequency was used for analysis. For SSM sorts, the ratio of mutant counts from a binding sort to the expression sort was compared to that of the wild type sequence, and the log difference was defined as an enrichment factor^46^.

### Synthesis of biotin conjugated small-molecules

Structures of small-molecule ligands conjugated to biotin are available in the supplementary materials.

### Expression and purification of designed small-molecule–binding proteins

DNA sequences of designed small-molecule binders were ordered as eblocks from Integrated DNA Technologies and cloned into an N-terminal hexahistidine tag-containing vector via the golden gate method as previously described^28^. The assembled plasmid was transformed into *E. coli* BL21(DE3) and resultant transformants were cultured in autoinduction media at 37 °C for 3-4 hours and 18 °C overnight. Cells were harvested by centrifugation at 4000 xg for 10 minutes. Cells were resuspended in lysis buffer (40 mM imidazole, 100 mM sodium phosphate, 500 mM sodium chloride, pH 7.4 or 40mM imidazole, 100mM Tris-HCl, and 500mM sodium chloride, pH 8.0) and then sonicated on ice for 5 minutes with 10 seconds on and 10 seconds off. Resulting lysates were centrifuged for 30 minutes at 14000 xg to clarify the lysate. The clarified lysates were applied to ∼1 mL of nickel resin and washed with 10 column volumes of wash buffer (40 mM imidazole, 100 mM sodium phosphate, and 500 mM sodium chloride, pH 7.4 or 30mM imidazole, 20mM Tris-HCl, and 500mM sodium chloride, pH 8.0). A pre-elution wash of 400 μL was performed with elution buffer (400 mM imidazole, 100 mM sodium phosphate, or 500 mM sodium chloride, pH 7.4 or 500mM imidazole, 20mM Tris-HCl, and 100mM NaCl, pH 8.0). Samples were eluted with 1.3 mL of elution buffer and then filtered through a 0.2 μm filter prior to SEC purification with an superdex 75 10/300 increase column. Final purified samples were snap-frozen in liquid nitrogen and stored at −80 °C.

### Characterization of binding affinity by ITC

Binding affinity of designed proteins was assessed by titrating purified protein with ligand using a microcal PEAQ ITC-Automated instrument. Both protein and ligand were prepared in identical buffers that were degassed by bottle-top filtration. Cortisol titrations contained 1% DMSO in all solutions and warfarin and apixaban titrations contained 5% DMSO in all solutions. All titrations were performed at 25 °C and 19 total injections, 0.4 μL for the first injection and 2 μL for the remaining 18. Resulting titration data were analyzed with the Malvern Panalytical ITC Analysis software, and a single-site binding model was used to fit the resulting data.

### Characterization of binding affinity by BLI

Biolayer interferometry (BLI) was used to estimate the binding affinity and screen for binding of the purified proteins and target small-molecules APX, and SN-38. HBS-EP+ buffer (Cytiva) with 0.1% Bovine Serum Albumin and 0.01% Tween20 was used to run the experiment and prepare the purified protein and functionalized ligand solutions. 1 μM of small-molecule targets with biotin conjugated were used as ligands to be immobilized to the Octet Streptavidin biosensors (Sartorius). Titration experiments were performed in 25C, and the kinetic response data was collected using Octet RED96 and R8 (ForteBio). The resulting responses of the analytes were corrected using the signal from the reference buffer, and further analysis was performed using the Octet Data Analysis software.

### Characterization of binding affinity by FP

Fluorescence polarization experiment was performed to determine the binding affinity of the OHP binder designs. The concentration of the fluorophore labeled version of 17α-hydroxyprogesterone, AlexaFluor488-PEG3-17α-hydroxyprogesterone (AF488-OHP) ^12^, was kept constant at 6 nM in phosphate buffered saline (PBS, Fisher BioReagents). The concentration of the stock fluorophore was measured by using the absorbance at 490 nm and the molar extinction coefficient 73,000 cm^-1^M^-1^. 2-fold serial dilutions of the binder protein were prepared in 24 wells with constant amounts of AF488-OHP, and the plate was incubated at room temperature, shaking for at least 30 minutes. Fluorescence polarization filter cube (Green FP, EX 485/20 nm;EM 528/20 nm, Agilent) was used to measure polarization with the plate reader (Synergy Neo2, Biotek). A binding isotherm model was used to fit the data and estimate K_D_.

### Surface redesign of hcy129 with ProteinMPNN

The design model of hcy129 was used as input to ProteinMPNN, and sequence design was performed with a temperature of 0.1 and cysteine was excluded during design. 10 sequences were generated and single-sequence structure predictions with AlphaFold were used to confirm that surface redesigns matched the original design model. Designs with Cα RMSDs less than 1 Å and plDDTs greater than 90 were manually inspected and designs that recapitulated the binding residue rotamers were selected for gene synthesis, biochemical characterization, and crystallography.

### Crystallography and structure determination of hcy129_mpnn5 bound to cortisol

For crystallography, hcy129_mpnn5 was expressed at 0.5 L volume in 2.8 L flasks in autoinduction media and incubated in a shaking flask at 250 rpm for 4-6 hours at 37 °C and then lowered to 18 °C and incubated overnight. Purification was performed as described above, except that the elution buffer for SEC was 100 mM CHES, 100 mM acetone oxime, 100 mM NaCl, pH 8.6. To the SEC eluate ([protein] ∼ 1 mg/mL), 0.5 M guanidinium hydrochloride was added and incubated for ∼10 minutes, after which NiCl_2_ was added to a final concentration of 1 mM and incubated overnight at room temperature for sequence-specific nickel-assisted tag cleavage (SNAC)^47^. After SNAC tag cleavage, the protein was applied to a nickel affinity column to remove the free tag as well as any uncleaved protein. Flow-through as well as 3–5 mL of wash buffer eluate were collected from the nickel column. A final SEC purification was performed in 20 mM HEPES, 50 mM NaCl pH 7.4 and the eluate was concentrated to 10 mg/ml and incubated with 1 mM cortisol prior to crystallization. This combined solution was used for crystal screening using a Mosquito LCP by STP Labtech. Crystals grew successfully in 0.2 M ammonium sulfate, 30% w/v PEG 8000 and were harvested directly from a screening tray, cryoprotected with 25% ethylene glycol, and stored in liquid nitrogen. X-ray diffraction was performed at APS 24ID-C, data were processed with XDS^48^, and the structure was phased by molecular replacement using the designed structure as the search model and refined with Phenix^49^.

### Computational design of cortisol-induced heterodimers

We employed the hcy129.1_CID-cortisol complex modeled using AlphaFold2 and GALiganddock as our target. This complex features three mutations in addition to hcy129.1, based on the SSM profile of hcy129 to serve as potential hydrophobic protein-protein interaction sites. The AlphaFold2 model of the triple-mutant (hcy129.1_CID) retains the backbone and pocket sidechain structure of hcy129.1. To design minibinders to the NTF2-cortisol interface, we first used PatchDock as previously described^26^ to find the initial seeding positions of the miniprotein scaffolds against the target interface, and subsequently created Rotamer interaction field (RIF) for both the exposed pocket residues of the NTF2 and the cortisol ligand. The miniprotein library described previously was docked using RIFdock to yield around 5 million total docks. A preliminary ‘predictor’ design step was used to rank the *in silico* designs using Rosetta ddG and contact molecular surface in which 1 million docks were selected for the downstream Rosetta design. The interface of the three components; minibinder, hcy129.1_CID (NTF2 protein), and cortisol, of selected docks were optimized by Rosetta FastDesign as described previously^26^ but with cortisol also being explicitly considered by Rosetta. All designs were filtered by contact molecular surface > 380, contact patch > 170, and Rosetta ddG < −35 prior to ProteinMPNN sequence redesign where residues within 5 Å of the cortisol ligand were fixed. Finally, we ran Alphafold2 prediction with the initial guess protocol^50^ and the designs passing pae_interaction < 10 and plDDT_binder > 85 were ordered as a synthetic oligo pool.

### Yeast display and FACS to screen for chemical induced dimerization

Yeast surface display library containing 60k designed minibinders was prepared as previously described^26^. After the induction of yeast cells in SGCAA medium supplemented with 0.2% glucose, cells were washed with PBSF and incubated with 1 μM purified biotinylated hcy129.1_CID, anti-c-Myc fluorescein isothiocyanate (anti-c-Myc-FITC, Immunology Consultants Laboratory) and streptavidin-phycoerythrin (SAPE, ThermoFisher) in the presence or absence of 1 μM cortisol for 1 hr at room temperature. Cell sorting was performed using a Sony SH800S cell sorter with software version 2.1.5. Three million cells were collected after applying the gate shown in **Fig. S10A** from the library that was incubated with 1uM cortisol. After recovery the collected cells were streaked on C-Trp-Ura agar plates. 96 colonies were randomly picked and were individually cultured in C-Trp-Ura media, followed by induction in SGCAA media. Yeast display and flow cytometry experiment was done with two conditions for each clone; 1. incubation with 0.2 μM biotinylated hcy129.1_CID, anti-c-Myc-FITC, and SAPE, and 2. Incubation with 0.2 μM biotinylated hcy129.1_CID, anti-c-Myc-FITC, SAPE, and 0.2 μM cortisol. All samples (96×2 conditions) were analyzed with the Invitrogen Attune flow cytometer after washing the samples in PBSF, and we observed the change in binding signal upon adding cortisol. 40 out of 96 clones exhibited substantial population shifts when comparing the two experimental groups. The 12 sequences out of 40 that showed the strongest cortisol-induced binding signal (**Fig. S10B**) were expressed in *E. coli* for further biochemical characterization.

### Characterization of cortisol-induced dimerization by SEC

An N-terminal AviTag construct of hcy129.1_CID and C-terminal his-tag-containing construct of mini11 as described above. The two protein components of the CID, Nterm-AviTag-hcy129.1_CID and miniH11-HHHHHH, were incubated at 1 μM in the presence or absence of 10 μM cortisol, incubated for ∼2 hours at room temperature, and injected onto an S75 increase 10/300 column with a running buffer of 20 mM HEPES, 50 mM NaCl, pH 7.4. Absorbance was monitored at 280 nm over the course of the elution and resulting elution profiles were overlaid to assess potential cortisol-induced shifts in elution.

### Characterization of cortisol-induced dimerization and cortisol sensing with NanoBiT fusions

hcy129.1_CID and miniH11 were genetically fused to SmBiT and LgBiT, respectively, and ordered as eBlocks from IDT. Synthetic genes were cloned into pET29b plasmid and transformed into BL21(DE3) and purified as described above. After purification of the CID sensor components, they were mixed together at 200nM each and then titrated with variable concentrations of cortisol and incubated for 2-3 hours, after which the luciferase substrate diphenylterazine (DTZ) was added at a final concentration of 25 μΜ. Immediately after adding DTZ, the luminescent signal was measured on a Neo2 plate reader. To estimate the K_D_ of the dimer by NanoBiT, the miniH11-LgBiT component was kept fixed at 0.1 μM and hcy129.1_CID-SmBiT titrated at variable concentrations.

## Contributions

G.R.L., S.J.P., C.N., and A.H-W.Y. conceptualized the project, developed the computational design pipelines, and experimentally characterized the designs. G.R.L., S.J.P., and C.N. contributed equally. D.B. supervised research. G.R.L., S.J.P., A.H-W.Y., and D.B. wrote the manuscript. D.T. and I.A. contributed to NTF2 scaffold design. J.D. provided LigandMPNN. J.A.M.M. performed the synthesis of biotinylated ligands under D.R.L.’s supervision. S.J.P., A.K., A.B., and H.N. performed crystallography experiments, data collection, and determined the crystal structure of the cortisol binder. I.G., D.V., and N.R. helped with yeast library assembly and deep sequencing. H.L.H. assisted with protein purification and BLI experiments. B.C. provided guidance on miniprotein design for the cortisol-induced dimerization system. H.K.H. helped with NGS data analysis.

## Acknowledgements

Crystallographic data was collected at the Advanced Photon Source (APS) Northeastern Collaborative Access Team beamline 24ID-C, which is funded by the National Institute of General Medical Sciences from the National Institutes of Health (P30 GM124165). This research used resources of the Advanced Photon Source, a U.S. Department of Energy (DOE) Office of Science User Facility operated for the DOE Office of Science by Argonne National Laboratory under Contract No. DE-AC02-06CH11357. We would like to thank the Wysocki lab at The Ohio State University for native mass spectrometry measurements of the cortisol CID system.

## Supplementary Figures

**Figure S1.**
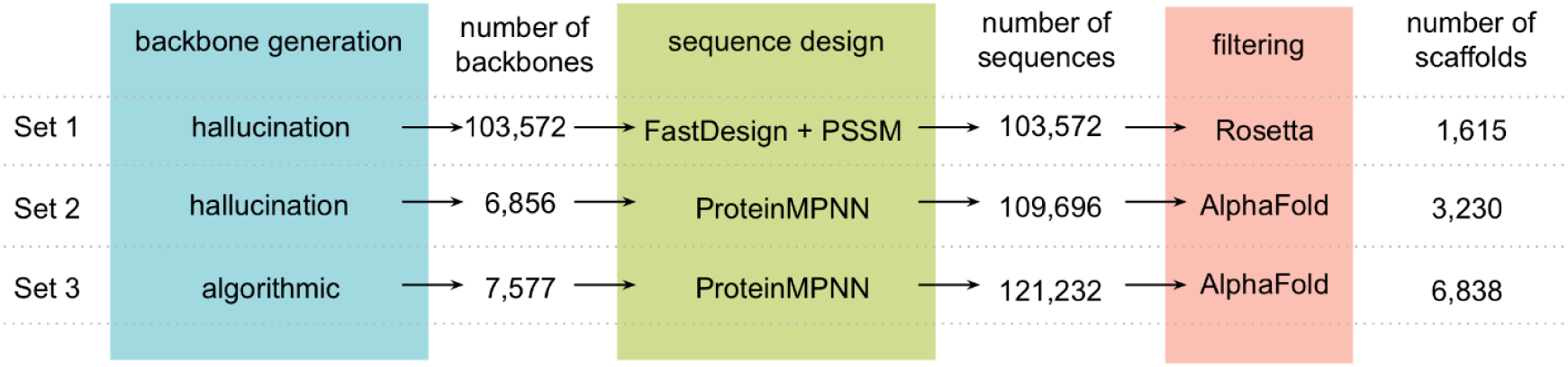
NTF2 scaffold generation. Computational pipelines to generate NTF2 scaffold sets.

**Figure S2.**
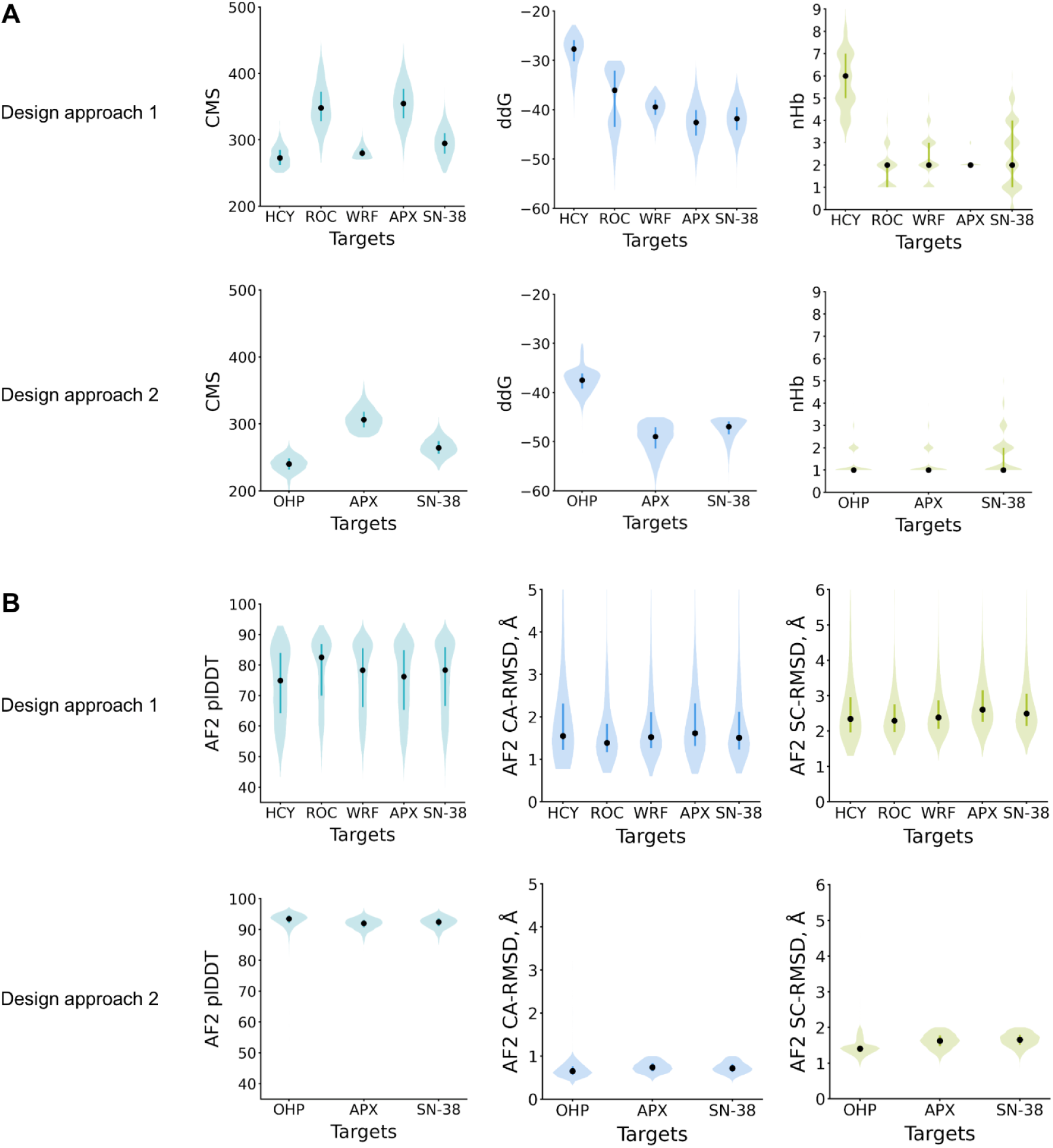
design metrics. (**A)** Rosetta metrics of ordered designs for each ligand that were used for selection; contact molecular surface (CMS), ddG, and number of protein-ligand hydrogen bonds (nHb). (B) AlphaFold metrics of selected designs for each small-molecule. CA-RMSD and the binding site sidechain RMSD (SC-RMSD) values are shown for the subset of designs with plDDT greater than 80.

**Figure S3.**
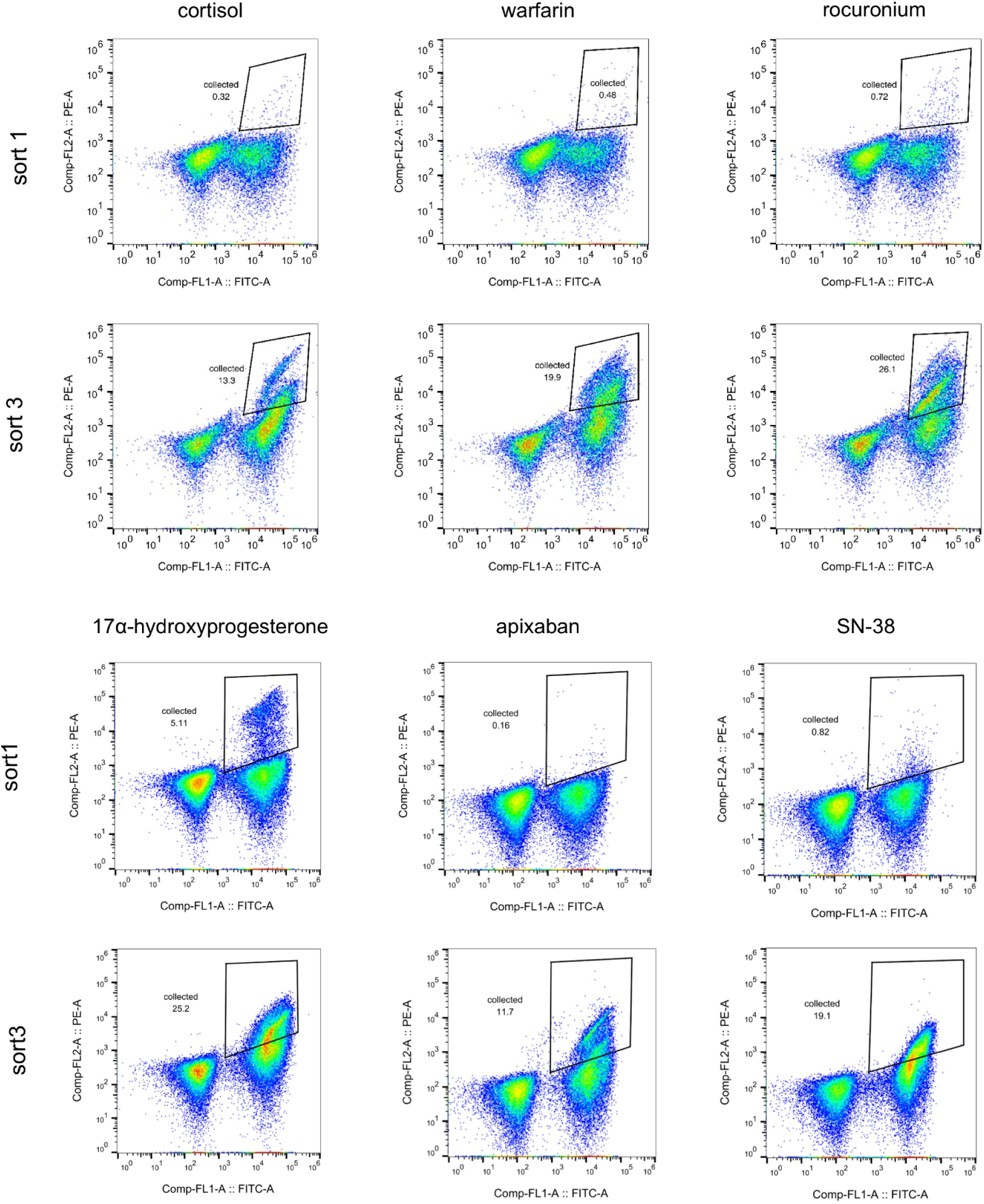
FACS plots of yeast display library screening. Representative FACS plots of designed protein yeast libraries. Libraries designed for cortisol, rocuronium, and warfarin were incubated with 1 μM of biotinylated small-molecule and 0.25 μM SAPE for all rounds. FACS results are shown for the 17α-hydroxyprogesterone (OHP) binder library incubated with 10 μM biotin-OHP without avidity (round1) and 2 nM without avidity (round3). For apixaban and SN-38 binder design libraries, FACS plots after incubation with 1 μM biotinylated ligand with avidity (round1), and 100 nM with avidity (round3) are shown.

**Figure S4.**
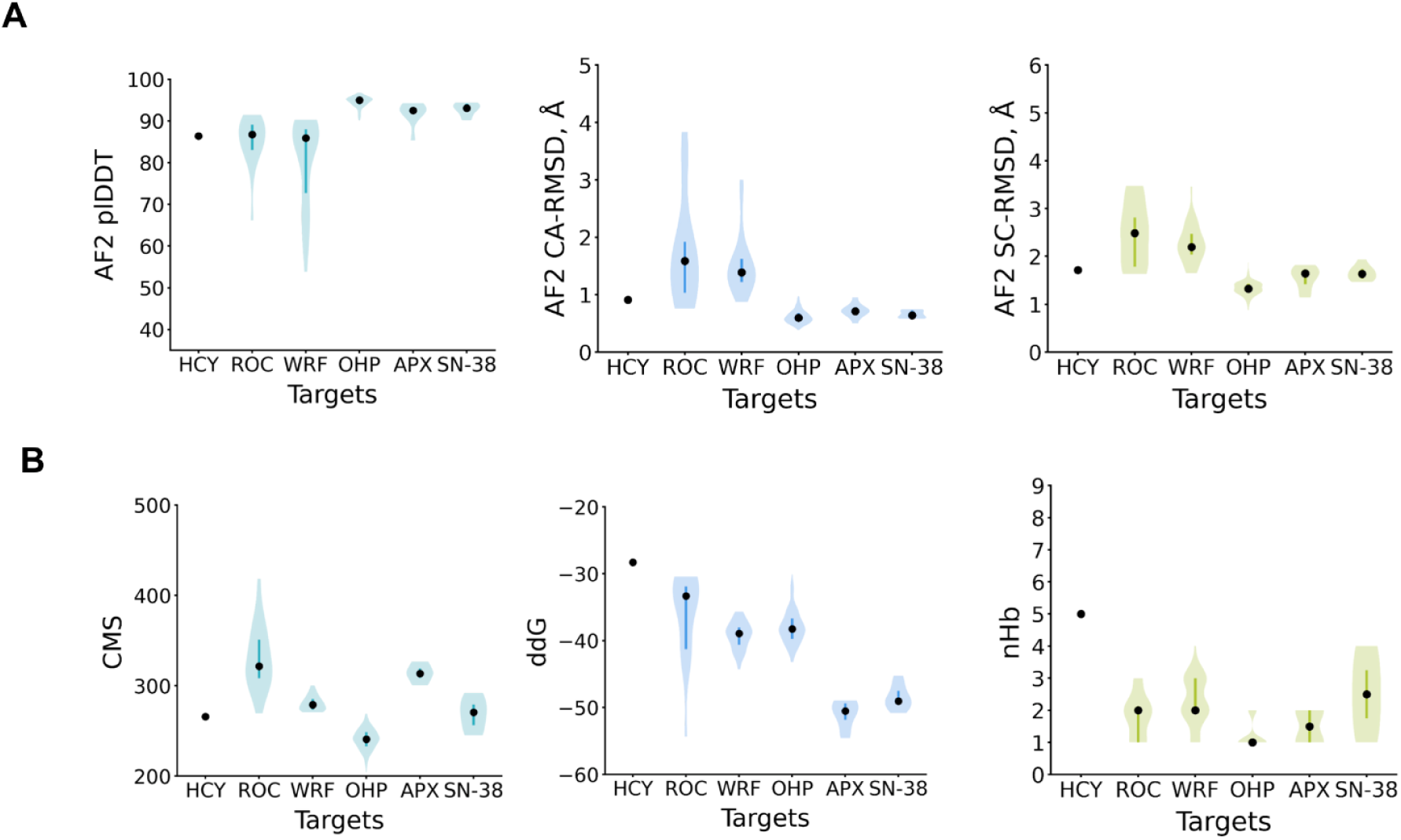
**(A)** AlphaFold2 (plDDT, Cα-RMSD of the models with plDDT > 80.0, and binding site side chain RMSD of the models with plDDT > 80.0) and **(B)** Rosetta metrics of putative binders identified by yeast display and FACS; contact molecular surface (CMS), ddG, and number of protein-ligand hydrogen bonds (nHb)

**Figure S5.**
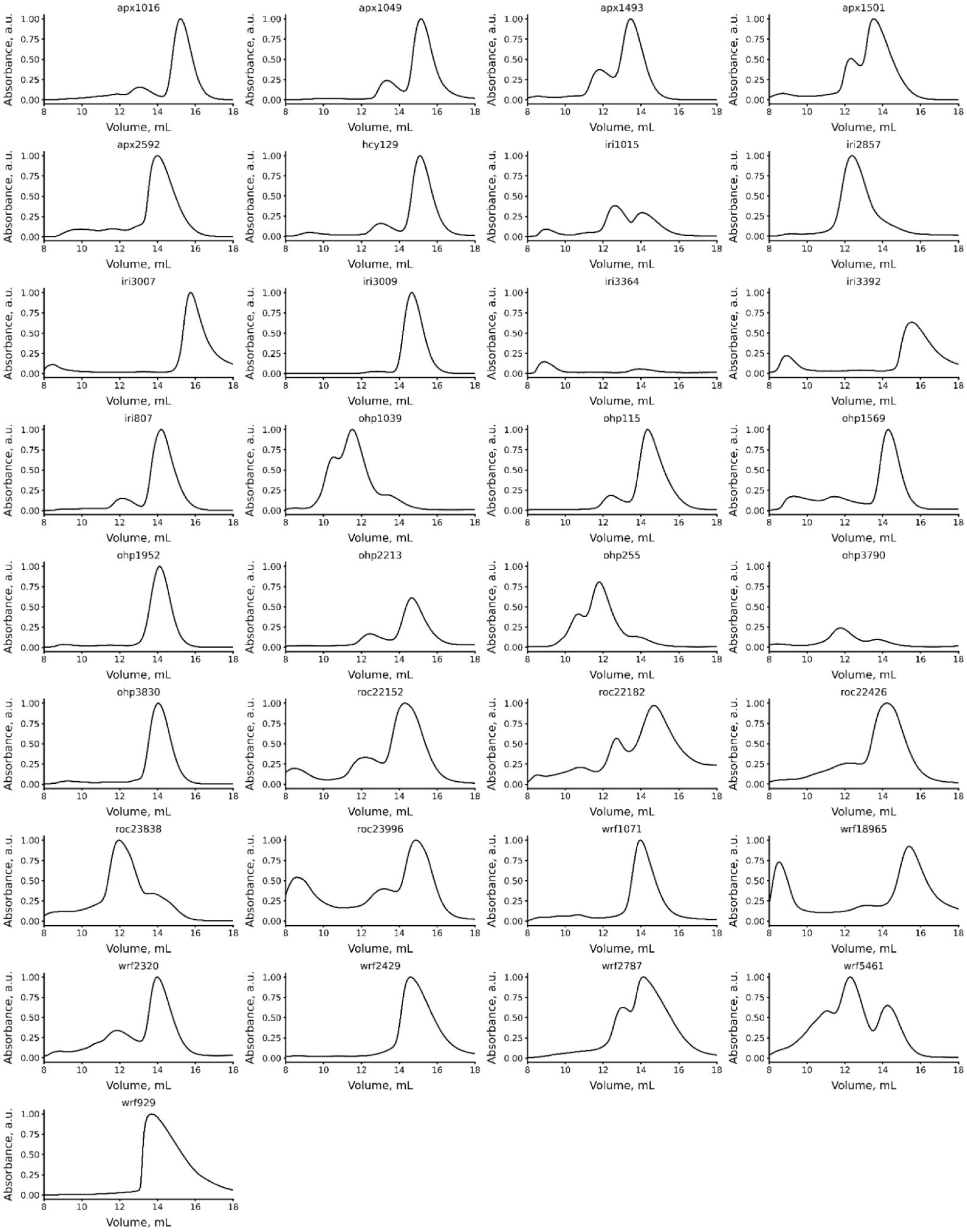
SEC elution profiles of nickel column eluates of putative binders identified from yeast display screening.

**Figure S6.**
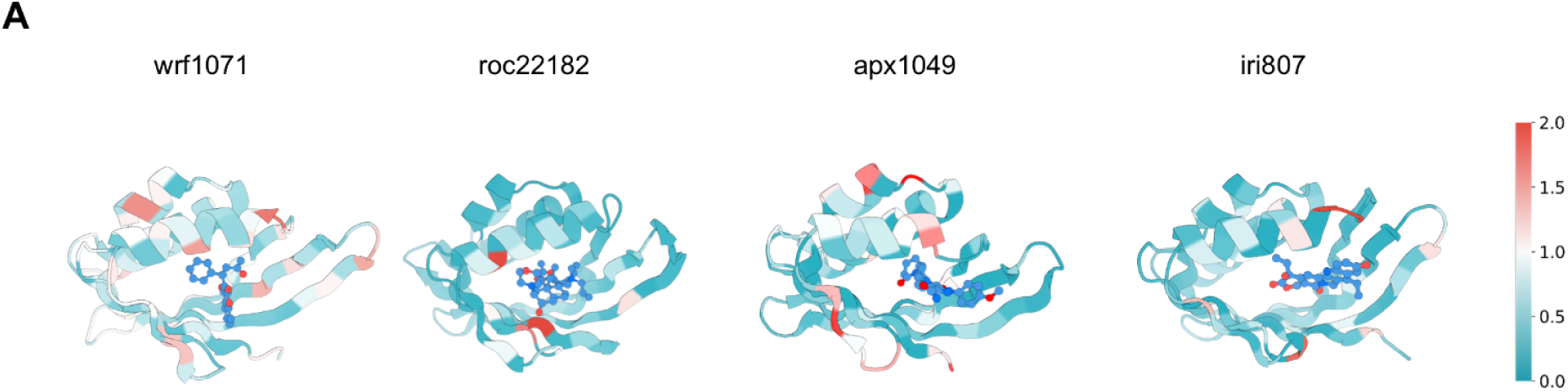
Site-saturation mutagenesis of select binder hits from yeast display. (**A**) The design models are colored by SSM sequence entropy (color bar heatmap teal: conserved, red: high entropy) for each position. Sequence entropy and binding enrichment analysis was done for wrf1071 SSM sort2 with 1uM biotin-warfarin without avidity, roc22182 SSM sort2 with 2uM biotin-rocuronium without avidity, apx1049 SSM sort2 with 1nM biotin-apixaban without avidity, and iri807 SSM sort2 with 10nM biotin-SN-38 without avidity.

**Figure S7.**
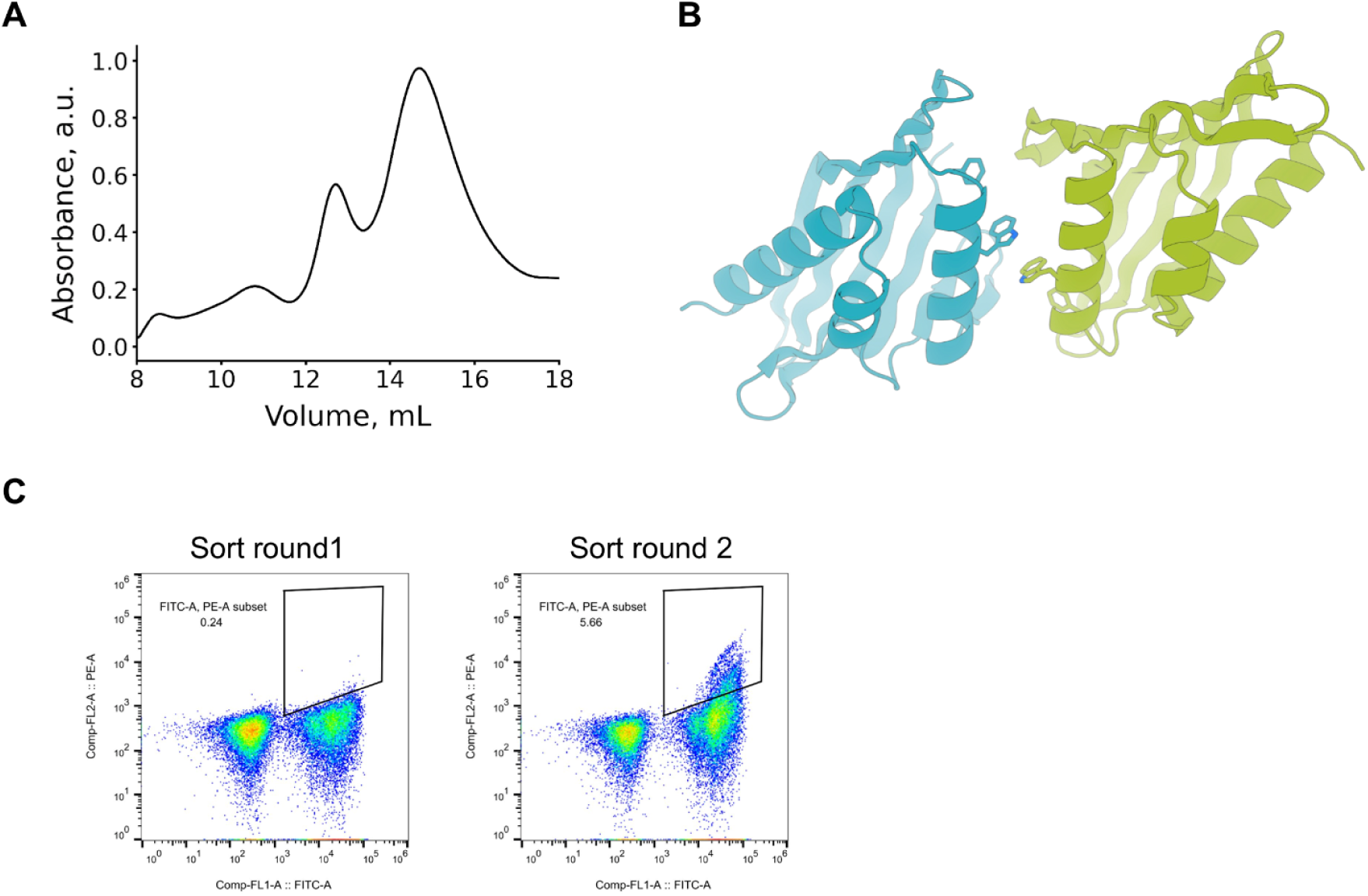
Structure-guided redesign of roc22182 to disrupt dimerization interface. (**A**) SEC trace of roc22182 sequence identified from yeast display FACS screening. (**B**) Dimer interface of roc22182 predicted by AlphaFold-multimer. (**C**) FACS plots of roc22182 combinatorial library incubated at 1 μM (round 1) and 100 nM (round 2) biotin-rocuronium without avidity.

**Figure S8.**
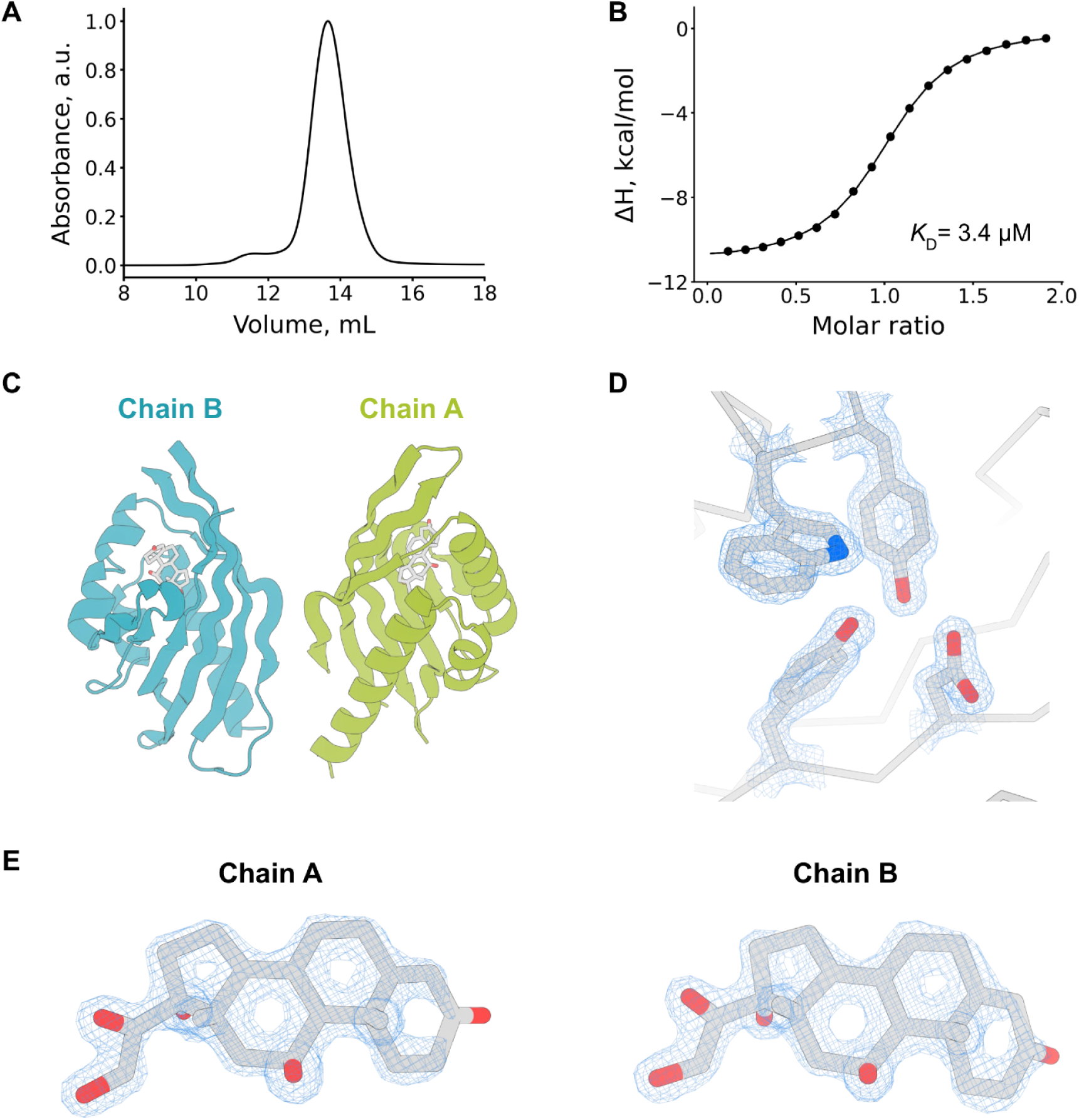
Biochemical and structural characterization of seq129_mpnn5. (**A**) SEC of MPNN-redesigned hcy129 variant hcy129_mpnn5. (**B**) ITC binding isotherm and raw heat for hcy129_mpnn5 titrated with cortisol (**C**) Asymmetric unit of hcy129_mpnn5 crystal structure (chain A: teal; chain B, green). (**D**) Representative 2Fo-Fc electron density of protein side chains in the hcy129_mpnn5 crystal structure contoured at 2σ. (**E**) 2Fo-Fc electron density of the ligand in Chain A and Chain B of the crystal structure contoured at 2σ.

**Figure S9.**
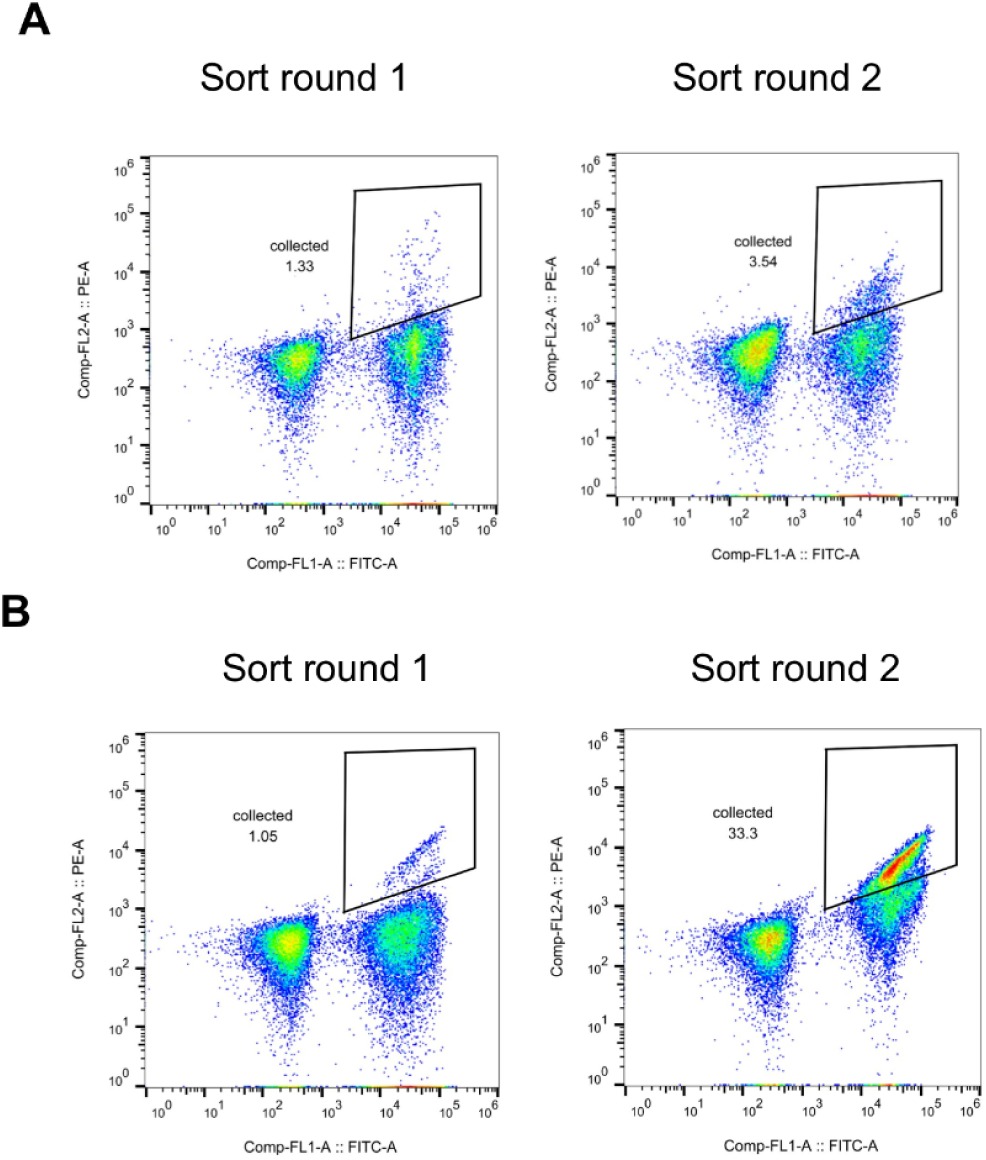
Optimization of the cortisol-binding protein seq129. (**A**) FACS plots of hcy129 SSM yeast library with sort 1 at 1 μM with avidity (left) and sort 2 at 100nM with avidity (right). (**B**) FACS plots of a hcy129 combinatorial mutant library containing favorable mutations identified from the SSM. Cells were collected for two consecutive rounds of FACS; for the first round the library was incubated with 100 pM biotin-cortisol without avidity, and the second round cell sorting was performed after incubation with 1 nM ligand without avidity.

**Figure S10.**
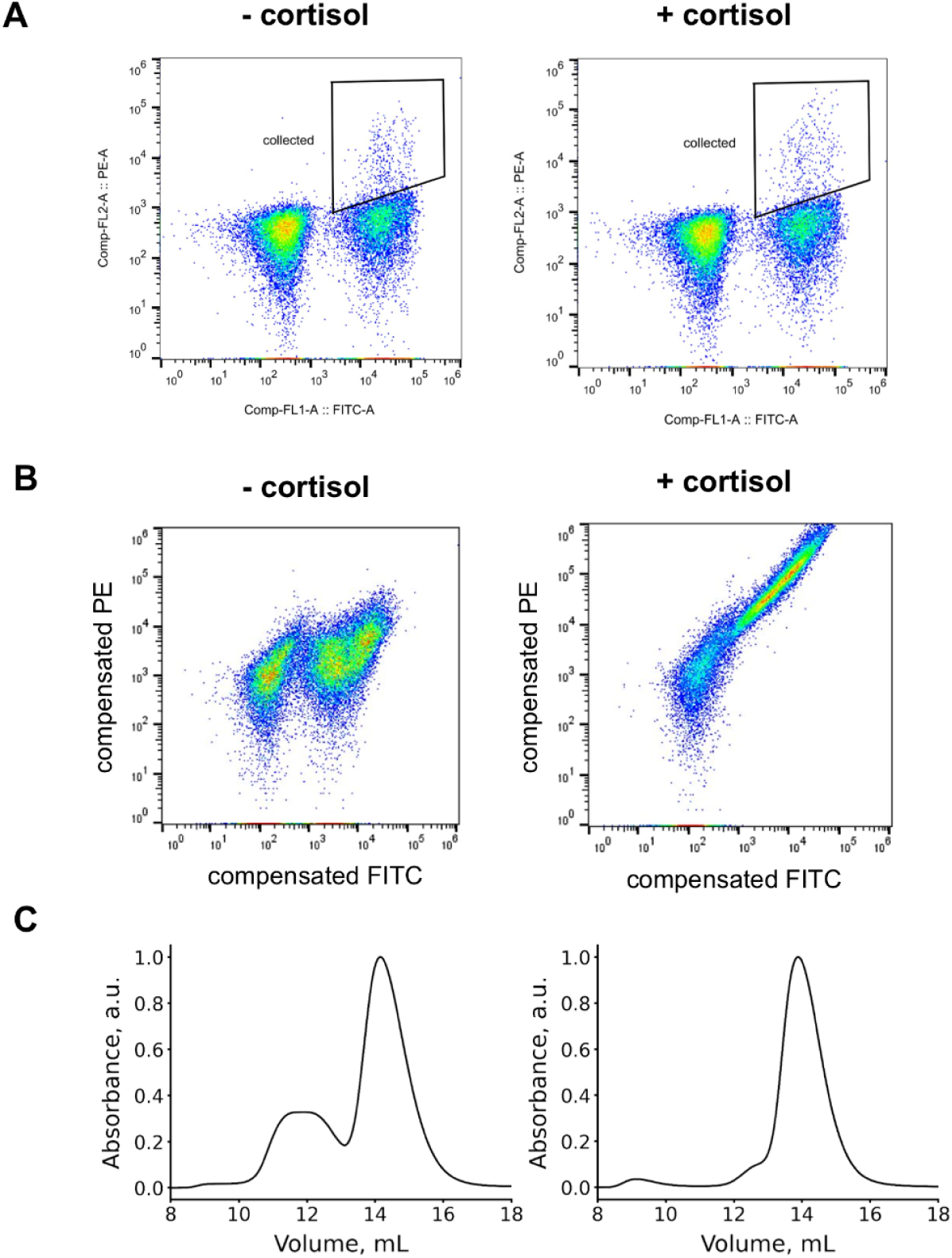
Library screening and hit characterization of designed CID minibinders. (**A**) FACS plots of yeast display library of minibinder CID library incubated with 1 μM biotinylated hcy129.1_CID in the absence (left) or presence (right) of 1 μM cortisol. (**B**) Representative single clone (miniH11) of yeast identified in libraries incubated with 0.2 μΜ biotinylated hcy129.1_CID in the presence (right) or absence (left) of 0.2 μΜ cortisol. (**C**) SEC traces of recombinant hcy129.1_CID (left) and miniH11 (right) proteins from *E. coli* expression.

**Figure S11.**
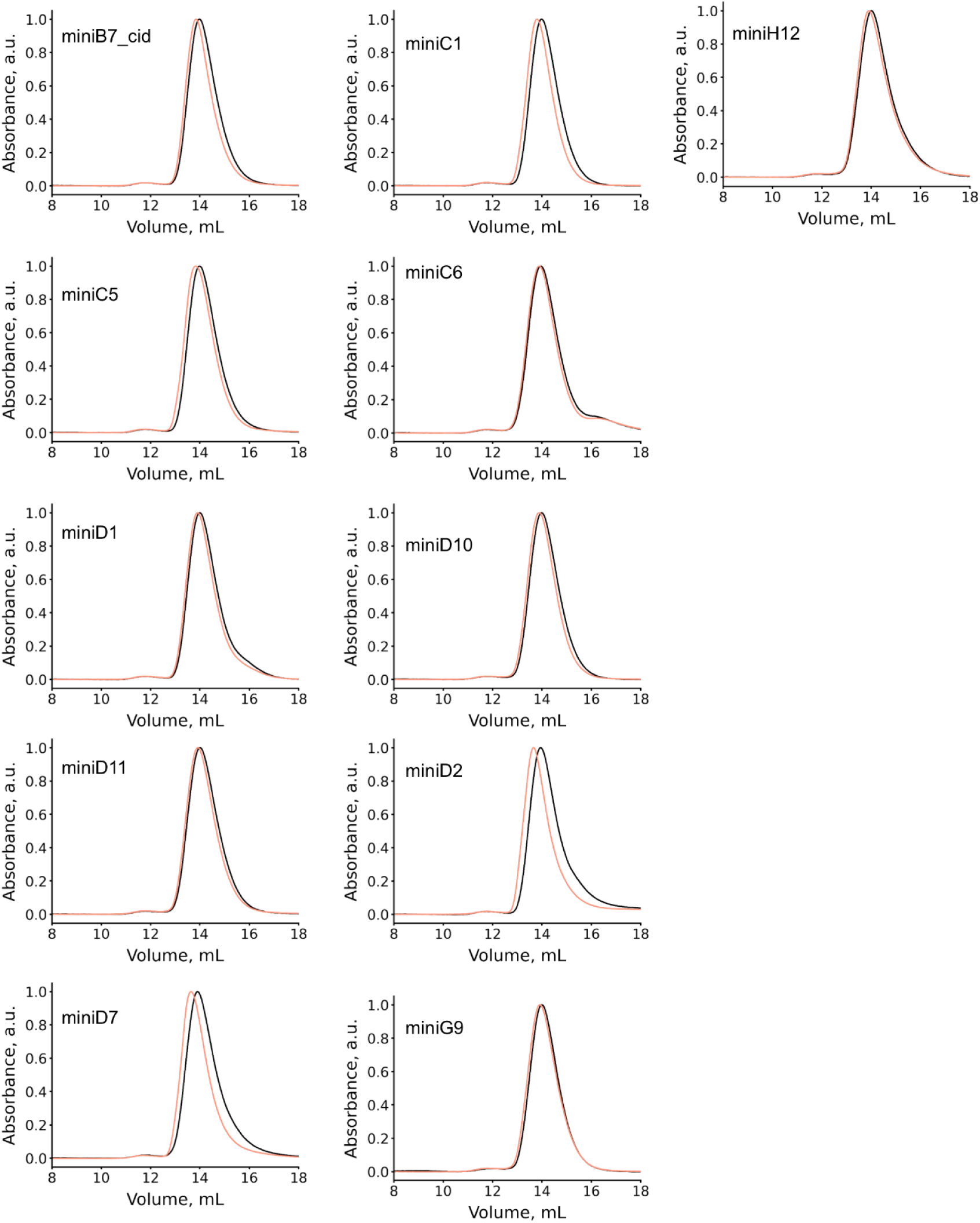
SEC of hcy129.1_CID (1 μM) combined with indicated miniproteins (1 μΜ) in the presence (peach) or absence (black) of cortisol (10 μM).

**Figure S12.**
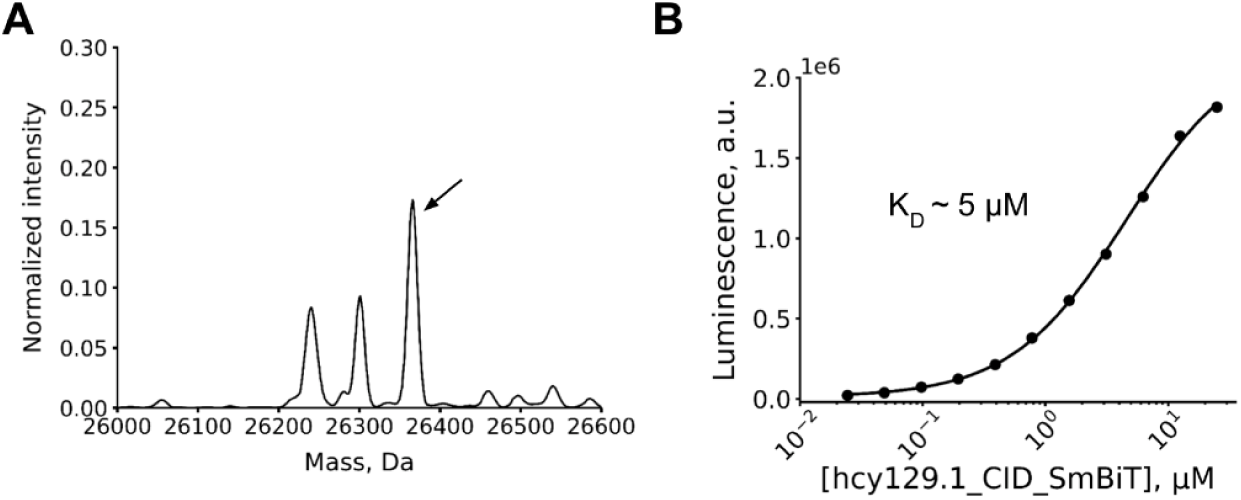
Characterization of mini11-seq129.1_CID cortisol-induced dimerization. (**A**) Native mass spectrum of mixture of hcy129.1_CID (1 μM), miniH11 (1 μΜ), and cortisol (10 μM) shows a distinct peak (indicated by black arrow) for the mass of the ternary complex at 26366 Da (expected: 26359 Da). (**B**) Binding isotherm of miniH11-LgBiT (100 nM) titrated with hcy129.1_CID-SmBiT.

## References

1. Stanton, B. Z., Chory, E. J. & Crabtree, G. R. Chemically induced proximity in biology and medicine. Science 359, (2018).

2. Glasgow, A. A. et al. Computational design of a modular protein sense-response system. Science 366, 1024–1028 (2019).

3. Taylor, N. D. et al. Engineering an allosteric transcription factor to respond to new ligands. Nat. Methods 13, 177–183 (2016).

4. Tinberg, C. E. et al. Computational design of ligand-binding proteins with high affinity and selectivity. Nature 501, 212–216 (2013).

5. Bick, M. J. et al. Computational design of environmental sensors for the potent opioid fentanyl. Elife 6, (2017).

6. Beltrán, J. et al. Rapid biosensor development using plant hormone receptors as reprogrammable scaffolds. Nat. Biotechnol. 40, 1855–1861 (2022).

7. Dou, J. et al. De novo design of a fluorescence-activating β-barrel. Nature 561, 485–491 (2018).

8. Polizzi, N. F. et al. De novo design of a hyperstable non-natural protein–ligand complex with sub-Å accuracy. Nat. Chem. 9, 1157–1164 (2017).

9. Polizzi, N. F. & DeGrado, W. F. A defined structural unit enables de novo design of small-molecule–binding proteins. Science 369, 1227–1233 (2020).

10. Basanta, B. et al. An enumerative algorithm for de novo design of proteins with diverse pocket structures. Proc. Natl. Acad. Sci. U. S. A. 117, 22135–22145 (2020).

11. Pan, X. & Kortemme, T. De novo protein fold families expand the designable ligand binding site space. PLoS Comput. Biol. 17, e1009620 (2021).

12. Dou, J. et al. Sampling and energy evaluation challenges in ligand binding protein design. Protein Sci. 26, 2426–2437 (2017).

13. Eberhardt, R. Y. et al. Filling out the structural map of the NTF2-like superfamily. BMC Bioinformatics 14, 327 (2013).

14. Bullock, T. L., Clarkson, D. W., Kent, H. M. & Stewart, M. The 1.6 Å Resolution Crystal Structure of Nuclear Transport Factor 2 (NTF2). J. Mol. Biol. 260, 422–431 (1996).

15. Yeh, A. H.-W. et al. De novo design of luciferases using deep learning. Nature 614, 774–780 (2023).

16. Dauparas, J. et al. Robust deep learning-based protein sequence design using ProteinMPNN. Science 378, 49–56 (2022).

17. Jumper, J. et al. Highly accurate protein structure prediction with AlphaFold. Nature 596, 583–589 (2021).

18. Hellhammer, D. H., Wüst, S. & Kudielka, B. M. Salivary cortisol as a biomarker in stress research. Psychoneuroendocrinology 34, 163–171 (2009).

19. Hanley, J. P. Warfarin reversal. J. Clin. Pathol. 57, 1132–1139 (2004).

20. Bom, A. et al. A novel concept of reversing neuromuscular block: chemical encapsulation of rocuronium bromide by a cyclodextrin-based synthetic host. Angew. Chem. Int. Ed Engl. 41, 266–270 (2002).

21. Lu, G. et al. A specific antidote for reversal of anticoagulation by direct and indirect inhibitors of coagulation factor Xa. Nat. Med. 19, 446–451 (2013).

22. Mathijssen, R. H. J. et al. Irinotecan pharmacokinetics-pharmacodynamics: the clinical relevance of prolonged exposure to SN-38. Br. J. Cancer 87, 144–150 (2002).

23. Morejón García, G., et al. Generation of monoclonal antibodies against 17α-hydroxyprogesterone for newborn screening of congenital adrenal hyperplasia. Clin. Chim. Acta 485, 311–315 (2018).

24. Boyken, S. E. et al. De novo design of protein homo-oligomers with modular hydrogen-bond network-mediated specificity. Science 352, 680–687 (2016).

25. Basanta, B. et al. Introduction of a polar core into the de novo designed protein Top7. Protein Sci. 25, 1299–1307 (2016).

26. Cao, L. et al. Design of protein-binding proteins from the target structure alone. Nature 605, 551–560 (2022).

27. Xiao, Y. & Woods, R. J. Protein-ligand CH-π interactions: Structural informatics, energy function development, and docking implementation. J. Chem. Theory Comput. 19, 5503–5515 (2023).

28. Wicky, B. I. M. et al. Hallucinating symmetric protein assemblies. Science 378, 56–61 (2022).

29. Laudat, M. H. et al. Salivary cortisol measurement: a practical approach to assess pituitary-adrenal function. J. Clin. Endocrinol. Metab. 66, 343–348 (1988).

30. Park, H., Zhou, G., Baek, M., Baker, D. & DiMaio, F. Force Field Optimization Guided by Small Molecule Crystal Lattice Data Enables Consistent Sub-Angstrom Protein–Ligand Docking. J. Chem. Theory Comput. 17, 2000–2010 (2021).

31. Dixon, A. S. et al. NanoLuc Complementation Reporter Optimized for Accurate Measurement of Protein Interactions in Cells. ACS Chem. Biol. 11, 400–408 (2016).

32. Landrum, G. RDKit: A software suite for cheminformatics, computational chemistry, and predictive modeling. http://www.rdkit.org/RDKit_Overview.pdf.

33. Wang, J., Wang, W., Kollman, P. A. & Case, D. A. Automatic atom type and bond type perception in molecular mechanical calculations. J. Mol. Graph. Model. 25, 247–260 (2006).

34. Bannwarth, C. et al. Extended tight-binding quantum chemistry methods. Wiley Interdiscip. Rev. Comput. Mol. Sci. 11, (2021).

35. Cole, J. C., Korb, O., McCabe, P., Read, M. G. & Taylor, R. Knowledge-Based Conformer Generation Using the Cambridge Structural Database. J. Chem. Inf. Model. 58, 615–629 (2018).

36. Tosco, P., Stiefl, N. & Landrum, G. Bringing the MMFF force field to the RDKit: implementation and validation. J. Cheminform. 6, 37 (2014).

37. Zubatyuk, R., Smith, J. S., Leszczynski, J. & Isayev, O. Accurate and transferable multitask prediction of chemical properties with an atoms-in-molecules neural network. Sci Adv 5, eaav6490 (2019).

38. Maguire, J. B. et al. Perturbing the energy landscape for improved packing during computational protein design. Proteins 89, 436–449 (2021).

39. Leaver-Fay, A. et al. ROSETTA3: an object-oriented software suite for the simulation and design of macromolecules. Methods Enzymol. 487, 545–574 (2011).

40. Tyka, M. D. et al. Alternate states of proteins revealed by detailed energy landscape mapping. J. Mol. Biol. 405, 607–618 (2011).

41. Klein, J. C. et al. Multiplex pairwise assembly of array-derived DNA oligonucleotides. Nucleic Acids Res. 44, e43 (2016).

42. Benatuil, L., Perez, J. M., Belk, J. & Hsieh, C.-M. An improved yeast transformation method for the generation of very large human antibody libraries. Protein Eng. Des. Sel. 23, 155–159 (2010).

43. Gietz, R. D. & Schiestl, R. H. Large-scale high-efficiency yeast transformation using the LiAc/SS carrier DNA/PEG method. Nat. Protoc. 2, 38–41 (2007).

44. Jacobs, T. M., Yumerefendi, H., Kuhlman, B. & Leaver-Fay, A. SwiftLib: rapid degenerate-codon-library optimization through dynamic programming. Nucleic Acids Res. 43, e34 (2015).

45. Zhang, J., Kobert, K., Flouri, T. & Stamatakis, A. PEAR: a fast and accurate Illumina Paired-End reAd mergeR. Bioinformatics 30, 614–620 (2014).

46. Rubin, A. F. et al. Correction to: A statistical framework for analyzing deep mutational scanning data. Genome Biol. 19, 17 (2018).

47. Dang, B. et al. SNAC-tag for sequence-specific chemical protein cleavage. Nat. Methods 16, 319–322 (2019).

48. Kabsch, W. XDS. Acta Crystallogr. D Biol. Crystallogr. 66, 125–132 (2010).

49. Liebschner, D. et al. Macromolecular structure determination using X-rays, neutrons and electrons: recent developments in Phenix. Acta Crystallogr D Struct Biol 75, 861–877 (2019).

50. Bennett, N. R. et al. Improving de novo protein binder design with deep learning. Nat. Commun. 14, 2625 (2023).

