## Supplementary materials for "Small-molecule binding and sensing with a designed protein family"

Gyu Rie Lee[1,2,3‡], Samuel J. Pellock[1,2‡], Christoffer Norn[1,2‡], Doug Tischer[1,2], Justas Dauparas[1,2], Ivan Anischenko[1,2], Jaron A. M. Mercer[4,5,6], Alex Kang[1,2], Asim Bera[1,2], Hannah Nguyen[1,2], Inna Goreshnik[1,2], Dionne Vafeados[1,2], Nicole Roullier[1,2], Hannah L. Han[1,2], Brian Coventry[1,2], Hugh K. Haddox[7], David R. Liu[4,5,6], Andy Hsien-Wei Yeh[8]*, David Baker[1,2,3]*
[1]Department of Biochemistry, University of Washington, Seattle, WA, USA.
[2]Institute for Protein Design, University of Washington, Seattle, WA, USA.
[3]Howard Hughes Medical Institute, University of Washington, Seattle, WA, USA.
[4]Merkin Institute of Transformative Technologies in Healthcare, Broad Institute of Harvard and MIT, Cambridge, MA, 02142, USA.
[5]Department of Chemistry and Chemical Biology, Harvard University, Cambridge, MA, 02138, USA.
[6]Howard Hughes Medical Institute, Harvard University, Cambridge, MA, 02138, USA.
[7]Division of Basic Sciences, Fred Hutchinson Cancer Center, Seattle, Washington, USA.
[8]Department of Biomolecular Engineering, University of California, Santa Cruz, CA, USA.

‡ Equal contribution

**Biotinylated small-molecules**
**ITC raw data and binding isotherms**
**FP binding isotherms**
**BLI sensograms**
**Supplementary Table 1.** Rosetta and AF2 metric cutoffs used to select designs
**Supplementary Table 2**. Small-molecule binding characterization results
**Supplementary Table 3.** Data collection and refinement statistics for the cortisol binder structure determination

### Biotinylated small-molecules used for FACS and BLI

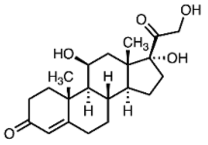

HCY

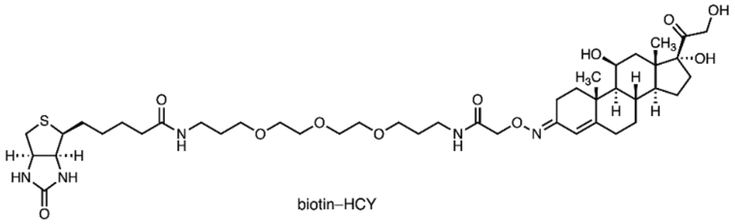

biotin–HCY

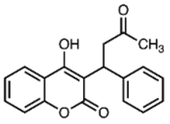

WRF

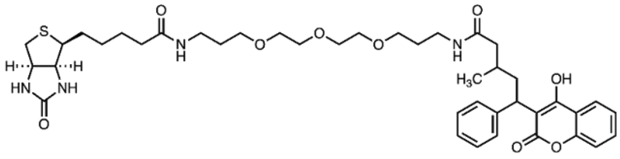

biotin–WRF

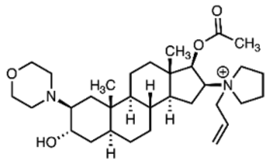

ROC

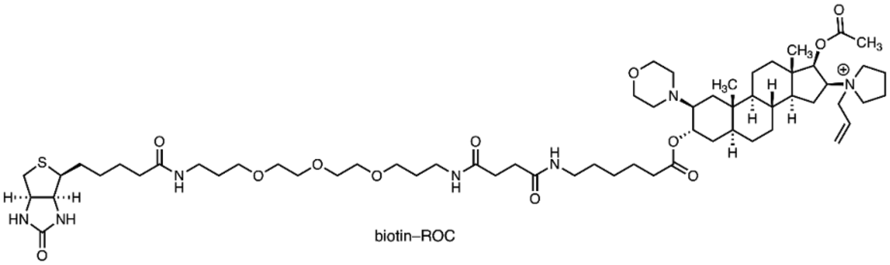

biotin–ROC

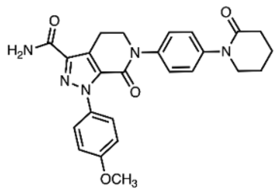

APX

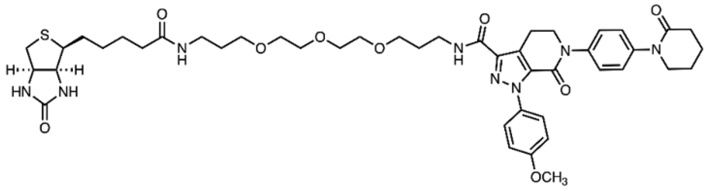

biotin–APX

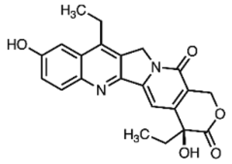

SN-38

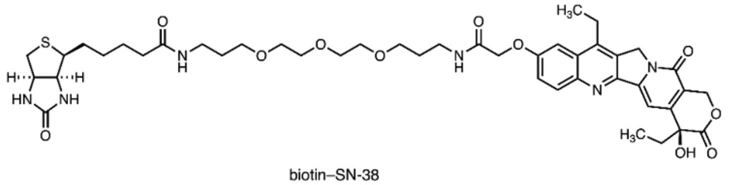

biotin–SN-38

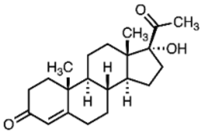

OHP

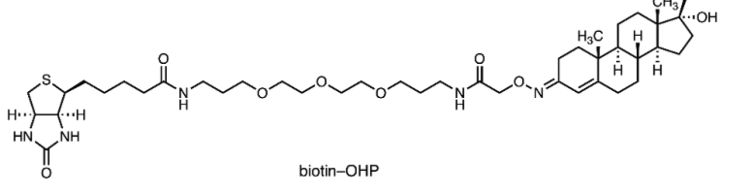

biotin–OHP

**ITC binding isotherm and corresponding raw data**

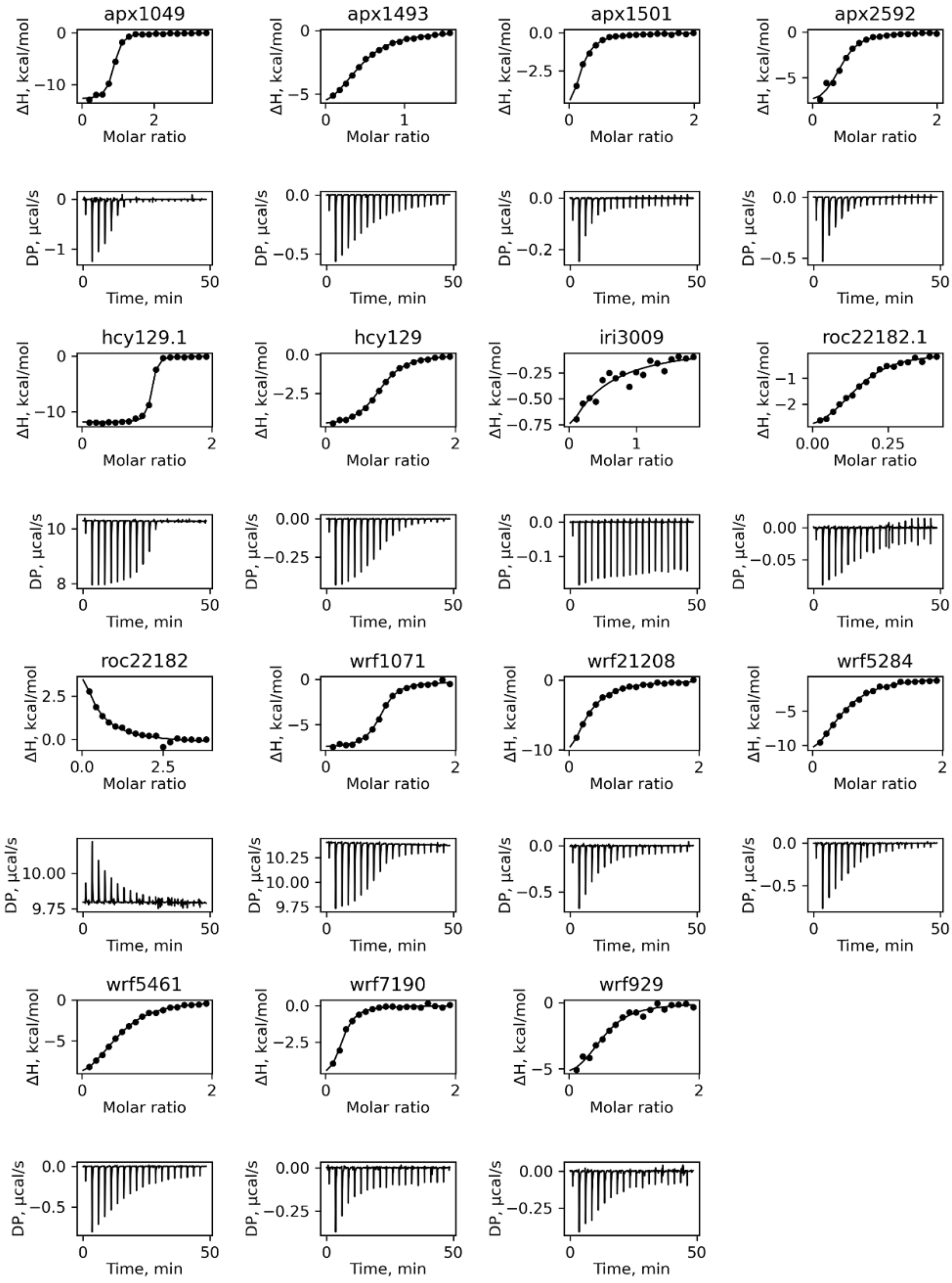

### FP binding isotherms

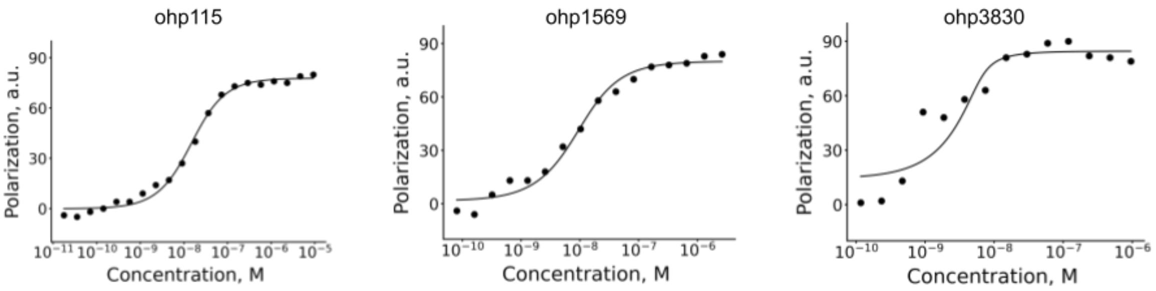


### BLI titration sensogram of SN-38 binder designs

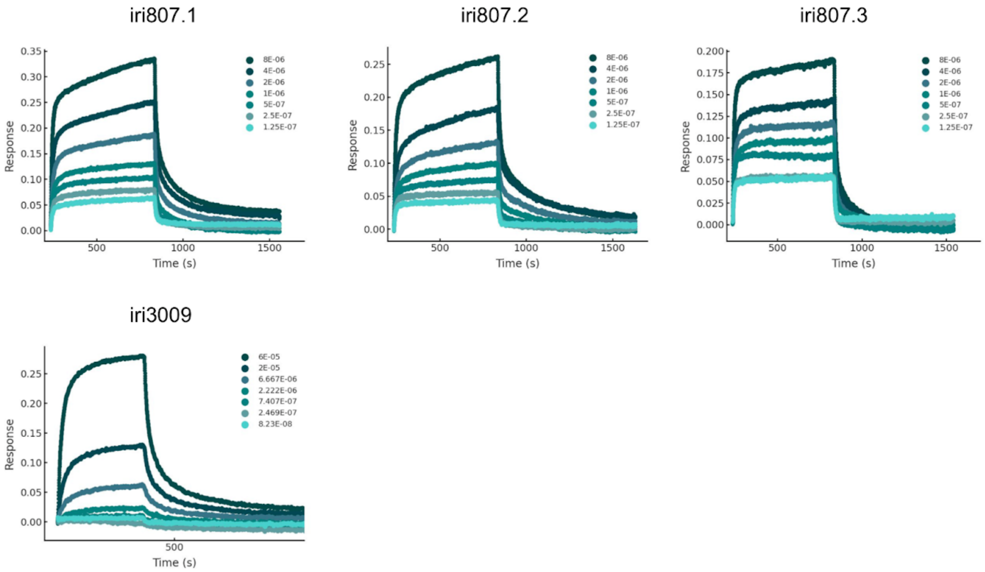

**Table S1**. Rosetta and AF2 metric cutoffs used to select designs

| Design approach | Target | Rosetta CMS | Interaction energy | Rosetta nHb | AF2 plDDT | AF2 Cα-RMSD (Å) | AF2 SC-RMSD (Å) |
|---|---|---|---|---|---|---|---|
| 1 | HCY | >250 | ddG/SASA<-0.09 | >1 | not applied | not applied | not applied |
| 1 | WRF | >270 | ddG/SASA<-0.13 | >1 | not applied | not applied | not applied |
| 1 | ROC | SC >0.65† | ddG<-30 | >1 | not applied | not applied | not applied |
| 1 | APX | >300 | ddG/SASA<-0.11 | >1 | not applied | not applied | not applied |
| 1 | SN-38 | >250 | ddG/SASA<-0.13 | >1 | not applied | not applied | not applied |
| 2-round1 | OHP | >200 | ddG<-35 | >=1 | >80 | <1 | <2 |
| 2-resample | OHP | >200 | ddG<-30 | >=1 | >80 | <1 | <1.5 |
| 2-round1 | APX | >250 | ddG<-40 | >=2 | >80 | <1 | <2 |
| 2-resample | APX | >280 | ddG<-45 | >=1 | >80 | <1 | <2 |
| 2-round1 | SN-38 | >230 | ddG<-40 | >=2 | >80 | <1 | <2 |
| 2-resample | SN-38 | >230 | ddG<-45 | >=1 | >85 | <1 | <2 |

†Used shape complementarity calculated by Rosetta instead of CMS

**Table S2**. Small-molecule binding characterization results

| **Target** | **Design name** | **Number of mutations compared to the original design** | **$K_D$ (μM)** | **$K_D$ determination method** |
|---|---|---|---|---|
| Cortisol | hcy129 | 0 | 2.3 | ITC |
| Cortisol | hcy129.1 | 4 | 0.065 | ITC |
| Cortisol | hcy129.1_CID | 7 | 0.064 | ITC |
| Coritsol | hcy129_mpnn5 | 9 | 3.4 | ITC |
| Warfarin | wrf1071 | 0 | 0.83 | ITC |
| Warfarin | wrf21208 | 0 | 10.9 | ITC |
| Warfarin | wrf5284 | 0 | 11.1 | ITC |
| Warfarin | wrf929 | 0 | 5.9 | ITC |
| Warfarin | wrf5461 | 0 | 8.8 | ITC |
| Warfarin | wrf7190 | 0 | 2.4 | ITC |
| Rocuronium | roc22182 | 0 | >50 | ITC |
| Rocuronium | roc22182.1 | 9 | 4 | ITC |
| Apixaban | apx1049 | 0 | 0.679 | ITC |
| Apixaban | apx1493 | 0 | 9.67 | ITC |
| Apixaban | apx1501 | 0 | 5.5 | ITC |
| Apixaban | apx2592 | 0 | 3.16 | ITC |
| 17α-hydroxy progesterone | ohp115 | 0 | 0.012 | FP |
| 17α-hydroxy progesterone | ohp3830 | 0 | <0.006 | FP |
| 17α-hydroxy progesterone | ohp1569 | 0 | 0.006 | FP |
| 17α-hydroxy progesterone | ohp1952 | 0 | 0.017 | FP |
| SN-38 | iri807 | 0 | n/a | BLI |
| SN-38 | iri807.1 | 3 | 0.77 | BLI (kinetic analysis) |

| | | | | |
|---|---|---|---|---|
| SN-38 | iri807.1 | 3 | 5.1 | BLI (Binding isotherm fitting using equilibrium response*) |
| SN-38 | iri807.2 | 3 | 1.0 | BLI (kinetic analysis) |
| SN-38 | iri807.3 | 3 | 5.47 | BLI (kinetic analysis) |
| SN-38 | iri3009 | 0 | 39 | BLI (kinetic analysis) |

*Dissociation could not be fit with kinetic analysis. $K_D$ of iri807.1 in **Figure 2D** is fit from max responses using a binding isotherm model.

**Table S3.** Data collection and refinement statistics for crystal structure of hcy129_mpnn5

| | **hcy129_mpnn5 (PDB: 8UQF)** |
|---|---|
| **Wavelength** | |
| **Resolution range** | 47.9 - 1.52 (1.57 - 1.52) |
| **Space group** | $P12_11$ |
| **Unit cell** | 48, 48, 56.4; 90, 94, 90 |
| **Total reflections** | 90862 (8830) |
| **Unique reflections** | 36533 (3628) |
| **Multiplicity** | 2.5 (2.4) |
| **Completeness (%)** | 90.3 (91.5) |
| **Mean I/sigma(I)** | 11.7 (1.4) |
| **Wilson B-factor** | 25.1 |
| **R-merge** | 0.06 (0.83) |
| **R-meas** | 0.08 (1.0) |
| **R-pim** | 0.04 (0.59) |
| **CC1/2** | 0.99 (0.61) |
| **CC*** | 0.99 (0.87) |
| **Reflections used in refinement** | 36181 (3627) |
| **Reflections used for R-free** | 1996 (195) |
| **R-work** | 0.23 (0.34) |
| **R-free** | 0.24 (0.35) |
| **CC(work)** | 0.94 (0.75) |
| **CC(free)** | 0.93 (0.66) |
| **Number of non-hydrogen atoms** | 2155 |
| **macromolecules** | 2022 |
| **ligands** | 57 |
| **solvent** | 76 |

| | |
|---|---|
| **Protein residues** | 246 |
| **RMS(bonds)** | 0.024 |
| **RMS(angles)** | 1.81 |
| **Ramachandran favored (%)** | 99.67 |
| **Ramachandran allowed (%)** | 0.4 |
| **Ramachandran outliers (%)** | 0.00 |
| **Rotamer outliers (%)** | 3.5 |
| **Clashscore** | 7.04 |
| **Average B-factor** | 33.1 |
| **macromolecules** | 33.0 |
| **ligands** | 29.3 |
| **solvent** | 39.9 |

Statistics for the highest-resolution shell are shown in parentheses.
